# Synaptic Activity Causes Minute-scale Changes in BAF Complex Composition and Function

**DOI:** 10.1101/2023.10.13.562244

**Authors:** S. Gourisankar, W. Wenderski, J. A. Paulo, S.H. Kim, K. Roepke, C. Ellis, S.P. Gygi, G.R. Crabtree

## Abstract

Genes encoding subunits of the SWI/SNF or BAF ATP-dependent chromatin remodeling complex are among the most enriched for deleterious *de novo* mutations in intellectual disabilities and autism spectrum disorder, but the causative molecular pathways are not fully known^1,2^. Synaptic activity in neurons is critical for learning and memory and proper neural development^3^. Neural activity prompts calcium influx and transcription within minutes, facilitated in the nucleus by various transcription factors (TFs) and chromatin modifiers^4^. While BAF is required for activity-dependent developmental processes such as dendritic outgrowth^5–7^, the immediate molecular consequences of neural activity on BAF complexes and their functions are unknown. Here we mapped minute-scale biochemical consequences of neural activity, modeled by membrane depolarization of embryonic mouse primary cortical neurons, on BAF complexes. We used acute chemical perturbations of BAF ATPase activity and kinase signaling to define the activity-dependent effects on BAF complexes and activity-dependent BAF functions. Our studies found that BAF complexes change in subunit composition and are selectively phosphorylated within 10 minutes of depolarization. Increased levels of the core PBAF subunit Baf200/*Arid2*, uniquely containing an RFX-like DNA-binding domain, are concurrent with ATPase-dependent opening of chromatin at RFX/X-box motifs. Changes in BAF composition and phosphorylation lead to the regulation of chromatin accessibility for critical neurogenesis TFs. These biochemical effects are a convergent phenomenon downstream of multiple growth factor signaling pathways in mouse neurons and fibroblasts suggesting that BAF integrates signaling information from the membrane. In support of such a membrane-to-nucleus signaling cascade, we also identified a BAF-interacting kinase, Dclk2, whose inhibition attenuates BAF phosphorylation selectively. Our findings support a direct role of BAF complexes in responding to synaptic activity to regulate TF binding and transcription.

## INTRODUCTION

ATP-dependent chromatin remodelers facilitate transcription factor (TF) binding and transcription by regulating the accessible chromatin landscape, by evicting, translocating, and remodeling nucleosomes^8^. Genes encoding remodelers are among the most highly mutated in neurodevelopmental disorders such as intellectual disabilities and autism spectrum disorder^1^. Subunits of the mammalian SWI/SNF complex, or BAF, are particularly enriched in *de novo* mutations in these disorders, implicating them as likely causative^1,2^. Recently, recessive mutations in neural-specific BAF complexes were shown to be causative of autism syndrome disorder^9^.

Synaptic activity in neurons leads to immediate (within minutes), *de novo* transcription of genes responsible for shaping social behavior, learning, and memory ^3^. Various chromatin modifiers, TFs, and remodelers, including BAF, have been identified as critical in facilitating the response to neural activity, membrane depolarization, and calcium influx ^4,10–14^. Signals from the synapse are integrated, in the nucleus, within minutes, often by phosphorylation of a TF or cytoplasmic-to-nuclear translocation of a factor, to guide transcription. However, the direct molecular consequences of synaptic activity on the BAF complex are not known.

BAF complexes are macromolecular machines assembled combinatorically from 15 different subunits encoded by 29 different genes^15,16^. Different protein interaction surfaces on subunits engender different interactions with TFs and different patterns of localization and activity on chromatin. At least three main classes of mammalian BAF complexes have been described that are present in all cells: canonical BAF (cBAF), polybromo-associated BAF (PBAF), and non-canonical BAF (ncBAF), each of which are defined by one of the two paralogous ATPases (BRG1 or BRM) and class-specific, non-redundant subunits^16,17^. For example, PBAF is defined by replacement of ARID1A/B (BAF250A/B) by ARID2 (BAF200) and BAF45B/C/D by PHF10 (BAF45A) as well as inclusion of PBRM1 (BAF180) and BRD7; it is larger by almost 0.5 MDa than cBAF^15,16^.

Homologs of subunits can also replace one another to encode cell-type- as well as functional specificity by their assemblies^15,18^. For example, during neural development, an exchange of neural progenitor-specific BAF subunits for neuron-specific subunits results in a neuronal BAF (nBAF) complex that is critical for neuronal function including activity-dependent processes such as dendritic arborization^5,6^. This complex is defined by the replacement of ACTL6A (BAF53A) by ACTL6B (BAF53B) and SS18 by SS181L (CREST) into canonical, polybromo-associated, or non-canonical BAF complexes^6,9,19^. While not all BAF complexes in neurons contain these subunits, their expression is sufficient for differentiation into neurons^20^.

We set out to map the membrane-to-nucleus pathway from synaptic activity to BAF complex function in neurons, modeling neural activity by an established protocol that depolarizes the membranes of primary mouse cortical neurons using KCl stimulation ^21,22^. Hypothesizing that activity prompts biochemical changes in BAF complexes, we conducted proteomic and phosphoproteomic studies that revealed a direct regulation of BAF combinatorial composition and stoichiometry within 10 minutes of membrane depolarization. Then, we mapped activity-dependent, BAF-regulated chromatin by using a potent (nanomolar IC_50_), specific cell-permeable inhibitor of BAF ATPase activity. We finally identified a select set of regulatory phosphorylations on BAF complexes that direct neural TF activity, and a potential BAF kinase, double-cortin-like kinase 2 (Dclk2). Our studies show at a minute-scale how BAF complexes are directly regulated at a molecular level by neural activity.

## RESULTS

### Activity-dependent switch in BAF complex composition

To examine the direct consequences on BAF complexes downstream of neural activity, we immunoprecipitated BAF from mouse primary cortical neuron cultures dissected from embryonic day 16.5 (E16.5) mice. The ATPase subunit Brg1, common to all BAF complexes, is approximately 2.2-fold more abundant in embryonic neuronal nuclear extract than its homolog, Brm, (Supplemental Table 1), so we immunoprecipitated BAF complexes with a monoclonal antibody specific to Brg1 (Fig. 1A). Neural activity was mimicked in culture by stimulation with 50 mM potassium chloride, an established protocol to depolarize membranes and stimulate calcium influx through L-type voltage-sensitive calcium channels (L-VSCCs) that leads to transcription of immediate-early genes (IEGs) such as *cFos* ^21,22^. Proteomic analysis identified 1,637 uniquely interacting proteins with BAF of which 1,452 had more than one peptide quantified, including each possible BAF subunit (Supplemental Table 2). Directly after stimulation, we observed a significant association of each of the four PBAF-specific subunits with Brg1 within just 10 mins of depolarization and increasing after 30 mins (Fig. 1B). Brg1 antibodies immunoprecipitated ∼50% more of Baf200/Arid2, Baf180/Pbrm1, Baf45a/Phf10, and Brd7 across 2-3 biological replicates (independent mice) than non-stimulated complexes (Fig. 1B). Immunoprecipitated subunits common to all BAF complexes such as Baf170/Smarcc2, Baf47/Smarcb1, or Baf57/Smarce1 stayed relatively constant except for a modest ∼20% decrease in Baf45b/Dpf1 at 10 mins and 1.2-fold increases in the ncBAF subunits Brd9, Bicra/Gltscr1, and Bicra/Gltscrl1 (Supplemental Fig. 1A). The total protein levels of BAF subunits did not change with depolarization, as measured by total nuclear extract mass spectrometry from the same cortical neurons used for Brg1 IP (Fig. 1C).

**Figure 1.**
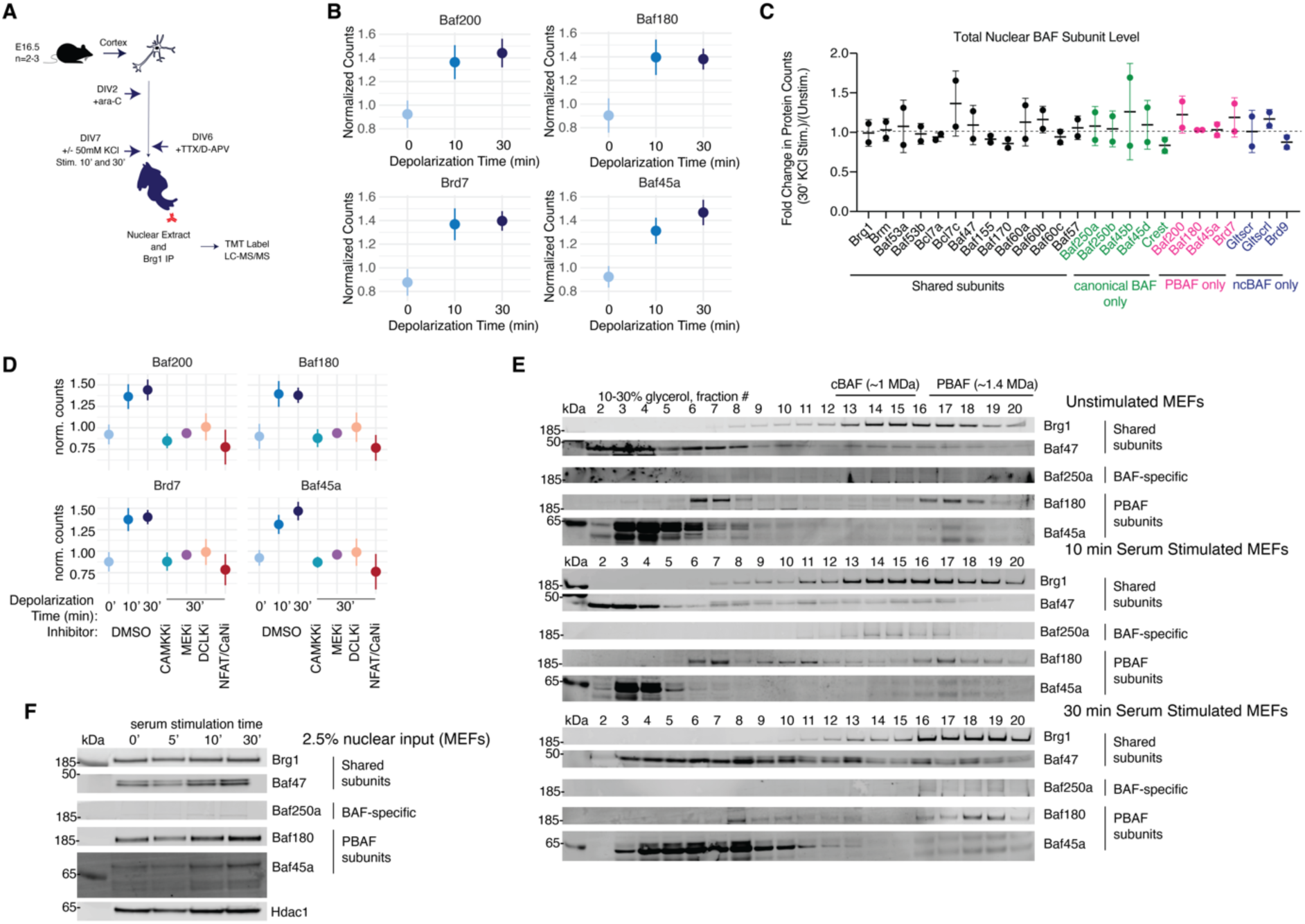
An activity-dependent switch in BAF complex composition. **A.** Schematic of immunoprecipitation-mass spectrometry (IP-MS) experiment to isolate cortical neuron BAF complexes. Anti-Brg1 antibody used for immunoprecipitation (Methods). **B.** Time-dependent increase in all four PBAF-specific subunits associated with Brg1 in cortical neurons, abundance normalized to Brg1 abundance; mean±s.d., n=2-3 biological replicates. **C.** Total levels of all BAF subunits in the nucleus of cortical neurons with time; mean±s.d., n=2 biological replicates **D.** Change in PBAF-specific subunits associated with Brg1 in cortical neurons after addition of kinase inhibitors to inhibit membrane signaling pathways; mean±s.d., n=2-3 biological replicates. CaMKKi: STO-609, 3 µM; MEKi: PD0325901, 3 µM; DCLKi: DCLK-IN-1, 2.5 µM; CaNi: 10 nM FK506 + 1 µM cyclosporin (CsA). Cyclosporin A and FK506 were used together because of the greater abundance of calcineurin in neurons compared to FKBP and cyclophilin, which are required for formation of inhibitor complexes^30^. **E.** Glycerol gradient fractionation of BAF complexes from stimulated mouse embryonic fibroblasts (MEFs), representative of n=2 biological replicates. **F.** Total levels of BAF subunits in MEF nuclear extracts, representative of n=2 biological replicates.

Calcium influx in neurons activates several membrane-to-nucleus signaling pathways, including mitogen-activated protein kinase (MAPK) signaling^23,24^, calcium/calmodulin-dependent kinase signaling (CaMK)^25–28^, and CaN (calcineurin)-NFAT (nuclear factor of activated T cell) signaling^29,30^. We treated neuronal cultures with nanomolar inhibitors to critical, exclusive kinases each of these signaling pathways and assessed BAF complex stoichiometry by Brg1- IP-MS (Supplemental Fig. 1B). Inhibiting each of these signaling pathways, on their own, reversed the switch in BAF complex composition within 30 minutes of drug addition and membrane depolarization. (Fig. 1D). There was little change in the relative levels of BAF subunits common to all complexes immunoprecipitated (Supplemental Fig. 1A).

Our results with kinase inhibitors in neurons (Fig. 1D) implied that the switch from cBAF to PBAF is a convergent effect of diverse membrane-to-nucleus signal cascades. To examine whether the switch in BAF complexes was a conserved phenomenon in multiple cell types, we fractionated BAF complexes in mouse embryonic fibroblasts (MEFs) that had been serum-starved and exposed to serum for 0, 10, or 30 mins (Fig. 1E). Quiescent fibroblasts respond to growth factors through MAPK signaling by rapidly inducing gene transcription^31^ in a similar manner to neurons^4,13^. Immunoblotting for Brg1, BAF, and PBAF complexes revealed a shift in Brg1 to heavier complexes at higher fractions (from fractions 13-16 to fractions 16-20) starting at 10 mins and increasing at 30 mins of serum exposure, representing Brg1-associated PBAF, which is ∼1.4 MDa compared to BAF at ∼1 MDa^16^. Total protein levels of BAF and PBAF subunits, normalized to Hdac1, remained similar (Fig. 1F). Hence the activity-dependent switch to PBAF, relative to the Brg1 ATPase, is a convergent, direct effect of membrane signaling in both neurons and fibroblasts.

### Identification of BAF-regulated, activity-dependent chromatin

To identify the direct targets of BAF downstream of neural activity, we profiled chromatin accessibility changes immediately (10 mins or 30 mins) after membrane depolarization with the addition of a 4 nM Brg1/Brm ATPase inhibitor, FHT1015 (Supplemental Fig. 2A,B). Changes in chromatin accessibility were measurable within 10 minutes of ATPase inhibition (Supplemental Fig. 2C), consistent with recent studies of acute BAF inhibition or degradation in mouse embryonic stem cells (mESCs)^32,33^. Of 23,902 peaks that changed statistically significantly (p_adj_≤0.05) and more than 2-fold after depolarization, almost 25%, or 5,944, were significantly different after BAF inhibition (p_adj_≤0.05) (Fig. 2B). Approximately half (2,710) of these BAF-regulated and activity-dependent peaks decreased in chromatin accessibility, almost all of which (93%) normally would have been upregulated after depolarization if there was no BAF inhibition (Fig. 2B, top). We therefore defined this set of peaks as the loci that required BAF ATPase activity for activity-dependent chromatin accessibility.

**Figure 2.**
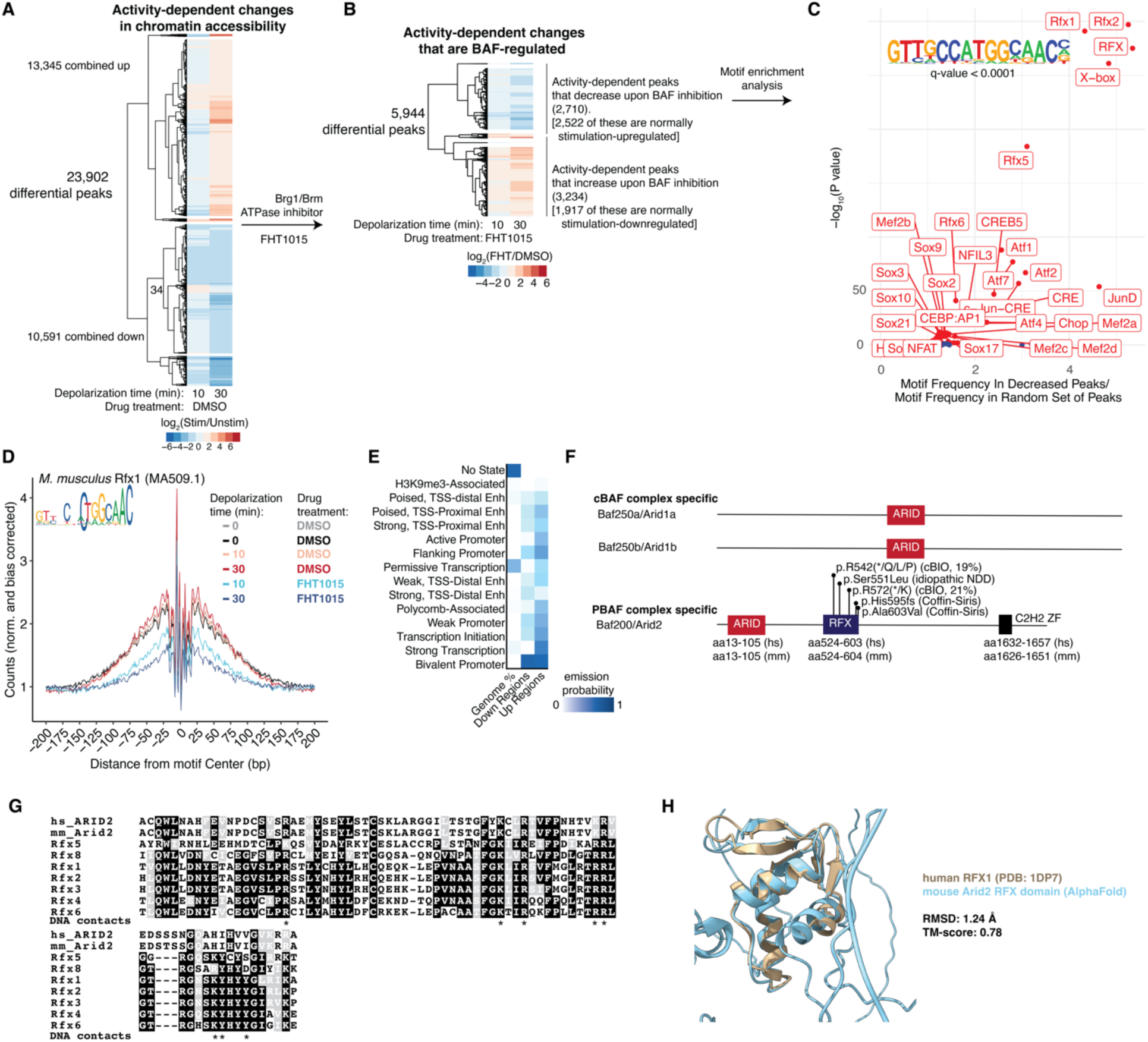
BAF regulates activity-dependent RFX accessibility. **A.** All activity-dependent changes in chromatin accessibility measured by ATAC-seq. **B.** Activity-dependent changes in chromatin accessibility that change statistically significantly upon simultaneous BAF ATPase inhibitor (FHT1015, 100 nM) treatment. For **A, B:** P-values computed by two-sided Wald test, adjusted for multiple hypotheses by Benjamini-Hochberg; differential peaks were classified as p_adj_≤0.05 and |log_2_(change)|≥ 1. **C.** Enrichment of transcription factor motifs in activity- dependent peaks that decrease upon BAF inhibition; P-values computed by two-sided Fishers’ exact test and q-value adjusted by Benjamini-Hochberg, labeled in red are q < 0.05. **D.** Footprint of chromatin accessibility at RFX motifs genome-wide. **E.** Enrichment of chromatin states in activity-dependent peaks that decrease or increase upon BAF inhibition from **B.** Chromatin state model trained on E16.5 mouse forebrain epigenome in^38^. **F.** Comparison of Arid subunits in canonical BAF (cBAF) and PBAF complexes, with mutations in neurodevelopmental disorders and cancers annotated for PBAF. **G.** Multiple sequence alignment of Arid2 and Rfx transcription factors in mouse and human with DNA contacts from RFX1 (PDB: 1DP7) annotated. **H.** Structural alignment of Arid2 Rfx domain and human RFX1.

These 2,710 sites that required BAF ATPase activity for activity-dependent accessibility were enriched (q-value (Benjamini-corrected) ≤ 0.0001) for the X-box motif, a 14-nucleotide imperfect repeat sequence bound by the evolutionarily conserved RFX transcription factors^34,35^ (Fig. 2C). Profiling accessibility at Rfx1 motifs genome-wide showed that depolarization stimulated a modest but significant increase in flanking accessibility, which was dramatically lost upon addition of the BAF ATPase inhibitor (Fig. 2D). The losses in accessibility spanned ∼300 bp centered on the motif (Fig. 2D). In a separate analysis to reproduce this finding, we used ChromVAR^36^ to quantify the variation in transcription factor motifs in all accessible peaks and found, again, that *M. musculus* Rfx1 (JASPAR MA0509.1) was the greatest (highest Z-score) driver of accessibility variation between inhibitor-treated and control datasets (Supplemental Fig. 2D). To characterize the chromatin landscape at these sites, we trained a hidden Markov model^37^ on the epigenome of an E16.5 mouse forebrain^38^. Most of the BAF-dependent, activity-dependent loci had the signature of bivalent promoters, characterized by H3K27me3 and H3K4me3 (Fig. 2E). Bivalent domains are often found at genes critical for differentiation and maintain these genes in a plastic state, ready to be induced upon receipt of a developmental signal^39^. These results suggest that BAF is involved in poising chromatin, at X-box sites, to respond to developmental signals.

The striking enrichment of X-box and RFX transcription factor motifs at BAF-required, activity-dependent loci made us investigate if RFX transcription factors interact biochemically with BAF complexes to direct their localization and activity. RFX transcription factors are evolutionarily conserved and known to direct the development of cilia on olfactory sensory neurons^40–42^. *De novo* deleterious mutations in human *RFX* genes have been linked to intellectual disability and autism spectrum disorder^43^. As neuronal BAF complexes have been shown to play critical roles in the development of olfaction circuitry in flies and in social behavior in mice and humans^9^, we hypothesized that Rfx transcription factors may interact with BAF in response to neural activity. However, in none of our proteomics or transcriptomic datasets, nor in public proteomic datasets, could we find any evidence to support such a direct interaction. Of the 8 mammalian RFX transcription factors (*RFX1-8*) in mammals, only 3 (Rfx3, Rfx1, and Rfx5) were detected in our total nuclear neuronal extracts by mass spectrometry (Supplemental Fig. 3A). Zero peptides of any of these were detected in our BAF IP-MS (Supplemental Fig. 3B). Only *Rfx3* and *Rfx7* were expressed at high levels in cortical neurons by RNA-seq (Supplemental Fig. 3C); however, neither those nor any of the other RFX transcription factors have been detected to interact with any BAF components in two different public databases of protein interactomes (STRING^44^ and BioGRID^45^) in either mouse or human cells.

### The BAF to PBAF switch can direct BAF localization and function

We noticed, instead, that the Baf200/Arid2 subunit specific to PBAF contains an RFX DNA-binding domain as well as a C2H2 zinc-finger (Fig. 2F). These domains are unique to Arid2 among the other BAF-complex Arids, Arid1a/Baf250a and Arid1b/Baf250b, which are both present only in canonical BAF complexes. The RFX domain in Arid2 is highly enriched in missense and truncation mutations in cancers^46^, and *de novo* mutations have also been found in idiopathic and Coffin-Siris Syndrome-like intellectual disabilities^47^ . We therefore speculated that the activity-dependent BAF-to-PBAF switch itself may guide BAF activity and localization on chromatin, via the introduction of RFX-DNA-binding capability in Arid2, a PBAF-specific subunit.

In support of this hypothesis are four observations. First, the Arid2 RFX domain is 100% identical between human and mouse and highly conserved with the winged-helix DNA-binding domains of mouse Rfx transcription factors (Fig. 2G). Almost all critical DNA-binding contacts are conserved in charge status; *i.e.* basic residues such as histidine, lysine, and arginine, are only replaced, if at all, by other basic residues (Fig. 2G). Second, a structural alignment of human RFX1^34^ with an AlphaFold model of the Arid2 RFX domain shows a very low root-mean-square deviation of ∼ 1.24 Å and a high template modeling score of 0.78 (1 indicates a perfect match, 0 indicates no match). Third, while none of the published cryo-EM structures of PBAF or its yeast homolog, RSC, have resolved the structure of the RFX domain in Arid2, density observed in a recent RSC cryo-EM structure^48^ indicated that it is flexibly attached to the structural core of the complex, and near extra-nucleosomal, linker DNA (Supplemental Fig. 3D). Finally, in yeast, RSC is known to bind both the nucleosome and the exit DNA, occupying wide, ∼ 150 to 300 bp nucleosome-depleted regions^49^. These contain both RSC and a partially-unwrapped nucleosome that is poised for removal and subsequent transcription^49^. Our analysis of BAF-regulated accessibility around RFX motifs showed that BAF ATPase inhibition reduces the flanking accessibility, spanning ∼ 300 bp, both in magnitude and in width, but not in footprint depth (Fig. 2D). This supports a model of PBAF binding to RFX sites and remodeling nucleosomes to generate chromatin accessibility, in response to developmental signals.

### Activity-dependent phosphorylation of BAF

Neural activity, as well as growth factor signaling and stimulation in other cell types, leads to activation of pre-existing, constitutively expressed transcription factors such as CREB (cyclic adenosine monophosphate (cAMP)-responsive element binding) and SRF (serum response factor) that drive immediate-early gene transcription^4^. The activation of these factors is by their post-translational modification, especially phosphorylation^50^. Careful functional investigation of their phosphosites has identified many of the regulatory kinase cascades that transduce membrane signals to these transcription factors, enabling a better understanding of how neural activity leads to transcription^25,27–30^. As BAF is also constitutively present in the nucleus, we hypothesized that neural activity leads to post-translational modification of BAF subunits that assist its regulation of chromatin accessibility.

Phosphoproteomic characterization of immunoprecipitated BAF complexes from cortical neurons revealed a specific set of ∼6 residues on 5 different BAF subunits significantly (p ≤ 0.05, fold change ≥ 1.5x) hyper-phosphorylated within just 10 minutes of depolarization (Fig. 3A). These likely represent direct targets of neural activity and represent the first identification of minute-scale activity-dependent phosphorylation of BAF complexes. Consistent with a recent report examining cortical neuron BAF phosphorylation after 1 h of depolarization^51^, serine 1349 on Brg1 (equivalent to serine 1382 in other Brg1 isoforms) was hyper-phosphorylated. After 30 minutes of depolarization, most of these same residues increased in phosphorylation, while a few residues on Brg1/Smarca4 and Dpf2/Baf45b became significantly hypo-phosphorylated (Fig. 3B). The changes were normalized to total immunoprecipitated protein abundance of these subunits, indicating that the phosphorylation detected was not due to the change in subunit abundance (if any). A total of 38 different phosphosites were identified on BAF subunits (Supplemental Table 2). Taken together, our data provides evidence of a biochemical kinase signaling pathway linking BAF with membrane depolarization.

**Figure 3.**
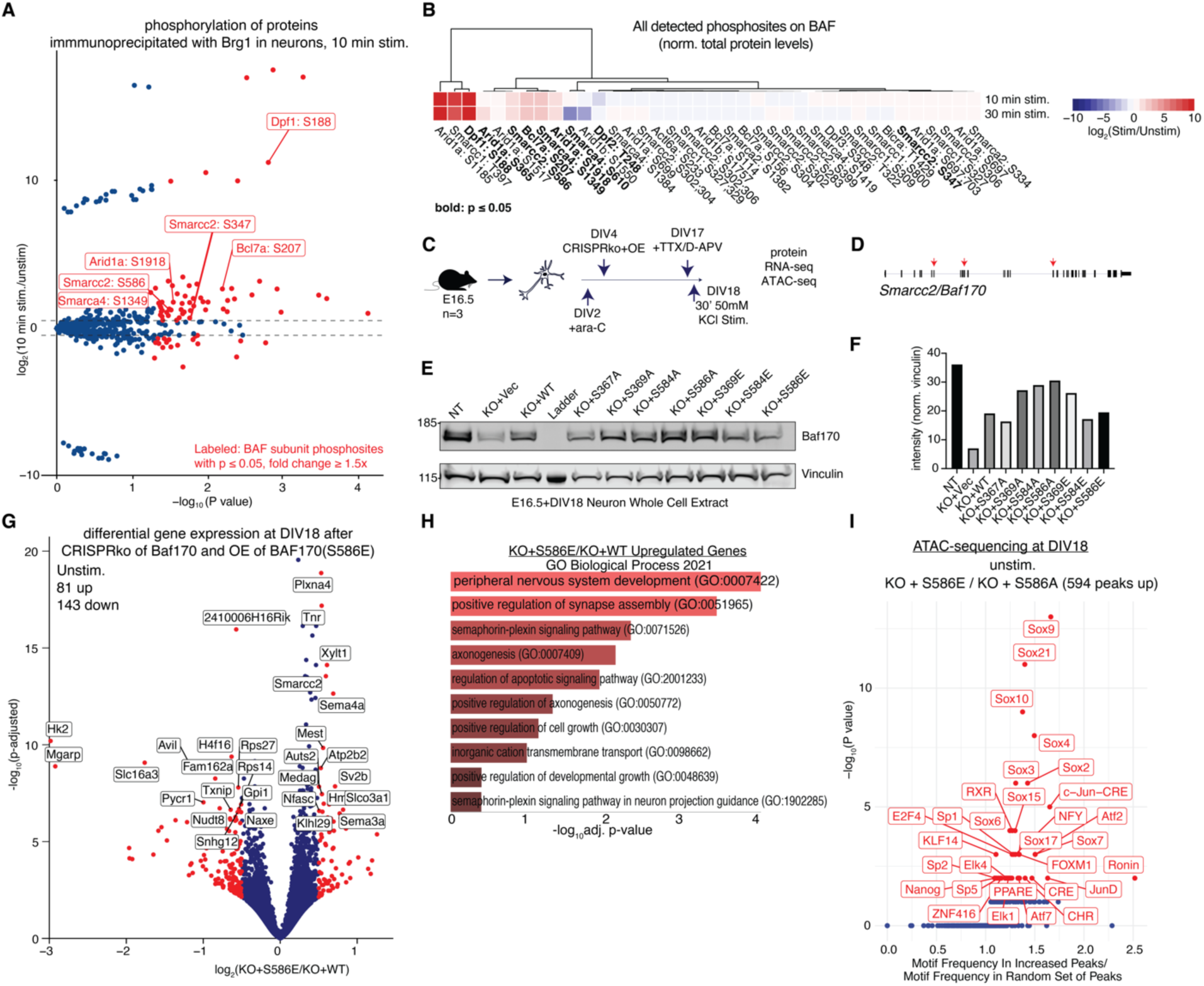
Activity-dependent phosphorylation of BAF complexes directs TF regulation. **A.** Differential phosphosites (norm. respective protein levels) detected after 10 min. stimulation of cortical neurons; P-values computed by two-sided Wald test, differential sites were classified as p_adj_≤0.05 and |log_2_(change)| ≥ 1.5. **B.** Changes in all detected phosphosites on BAF subunits only as a function of stimulation time. **C.** Schematic for knockout and add-back experiment to study Baf170 phosphosites. **D.** gRNA targeting sites on Baf170 gene. **E.** Total protein levels of overexpressed mutants and knockout. **F.** Quantification of **E. G.** Volcano plot of changes in unstimulated DIV18 cortical neurons between those that had Baf170KO and Baf170(S586E) and those that had Baf170KO and Baf170 (wild-type, WT) added back; n=2-3 biological replicates. P-values computed by two-sided Wald test, adjusted for multiple hypotheses by Benjamini-Hochberg; differential genes were classified as p_adj_≤0.05 and |log_2_(change)|≥ 0.5. **H.** GO Biological Processes enriched in upregulated genes in **G**; P-values computed by Fisher’s exact test corrected for multiple hypotheses by Benjamini-Hochberg. **I.** Enrichment of transcription factor motifs in increased chromatin accessibility peaks in DIV18 neurons that change between Baf170 KO + S586 status; P-values computed by two-sided Fishers’ exact test and q-value adjusted by Benjamini-Hochberg, labeled in red are q < 0.05; n=2 biological replicates.

### Functional characterization of activity-dependent Baf170 phosphorylation

To identify the possible regulatory consequences of BAF phosphorylation, we focused on the core subunit Baf170/Smarcc2 which was hyper-phosphorylated after depolarization at serine 586 in multiple of our BAF IP-MS and nuclear extract MS datasets. Phosphosites on Baf170 have also been characterized to regulate neurogenesis^52^ and Baf170/Smarcc2 is required for the formation of neural progenitors from embryonic stem cells^53^. *SMARCC2,* the human gene encoding BAF170, is also a high-confidence autism spectrum disorder gene^54^ and deleterious *de novo* variants have been found in individuals with neurodevelopmental delay^55^, suggesting that regulation of BAF170 may play a rate-limiting role in the processes of neural development.

We generated Baf170 knockout cortical neurons in culture and replaced the knockout with wild-type Baf170 or un-phosphorylatable (serine-to-alanine) and phospho-mimic (serine-to-glutamate) Baf170 by overexpression (Fig. 3C,D). Knockout was ∼80% complete after 18 days of culture (DIV18) and mutants were well-expressed at levels comparable to wild-type (Fig. 3E,F). As the Baf170(S586) residue was significantly and immediately hyper-phosphorylated after depolarization, we first assayed the consequences of mutating it to an un-phosphorylatable S586A (KO+S586A). To our surprise, RNA-seq analyses in neurons after 30 minutes of depolarization showed negligible changes in gene expression, as compared to neurons with Baf170 knockout and addback of a wild-type Baf170 copy (KO+WT) (Fig. 3G,H). Principal component analysis showed that the transcriptome of the KO+S586A mutant was most similar to the KO+WT neurons (Fig. 3G), and only 19 genes were differentially regulated (13 up, 6 down, p_adj_≤0.05, |log_2_(Fold Change)|≥0.5). In an orthogonal assay at a shorter time point in neuronal culture (DIV4), we looked by microscopy at activity-dependent dendritic outgrowth, a known response of neurons to depolarization that depends on BAF complexes^5–7^. As expected, Baf170 knockout reduced activity-dependent growth, and the addback of the wild-type Baf170 rescued this defect (Fig. 3I,J). Consistent with the transcriptomic finding, however, addback of the S586A mutant also rescued the defect in activity-dependent outgrowth (Fig. 3I,J). Therefore, we concluded that the single hyper-phosphorylation on S586 was not essential for activity-dependent processes.

### Baf170 phosphorylation directs Sox TF activity

In our transcriptomic data, we noticed that the S586E addback clustered close to the Baf170 knockout transcriptome in resting (unstimulated) neurons (Fig. 3G). This mutant was nevertheless well-expressed in cells (Fig. 3E). Comparing this mutation (KO+S586E) with wild-type Baf170 addback (KO+WT) across the transcriptome, we found 224 genes differentially regulated (81 up, 143 down) (Supplemental Fig. 4C). Upregulated genes were enriched for biological processes related to nervous system development (p_adj_≤0.0004) and synapse assembly (p_adj_≤0.0004) (Supplemental Fig. 4D).

Hypothesizing that constitutive phosphorylation, mimicked by the S586E mutant may introduce a neo-interaction that directs BAF-regulated chromatin accessibility at loci important in neural development, we conducted ATAC-sequencing in resting neurons with the S586E and S586A mutants, and analyzed differential motif accessibility. While only a few sites (38 peaks) had decreased accessibility in KO+S586E compared to KO+S586A (Fig. 3K), 594 loci had increased accessibility (Fig. 3L). These were enriched for Sox transcription factor motifs, especially Sox9, Sox21, and Sox10 (p≤10^-8^) (Fig. 3L). Each of these and other Sox transcription factors direct different neural differentiation pathways: Sox9 is important in the generation of astrocytes; Sox21 for neurons; Sox10 for oligodendrocytes^56^. Sox motifs were not enriched in differential peaks caused by Baf170 knockout (compared to wild-type), suggesting that the effect on Sox TF accessibility was uniquely due to introduction of a phosphosite-mutated Baf170 (Supplemental Fig. 4E). Our data therefore nominates the hyper-phosphorylation of Baf170 as helping BAF regulate accessibility for the activity of neurogenic transcription factors.

### Identification of Dclk as a putative BAF kinase

Our phosphoproteomic data on BAF complexes revealed a membrane-to-nucleus biochemical pathway resulting in phosphorylation of BAF subunits, and thus we sought to nominate a BAF kinase. In our BAF IP-MS data, we noticed that one of the proteins significantly (p_adj_ ≤ 0.05) increased in association with BAF after 10 minutes of depolarization was Dclk2 (doublecortin like kinase 2) (Fig. 4A). Dclk2 and its close homolog, Dclk1, are kinases almost exclusively expressed in the brain and contain both doublecortin domains and a C-terminal serine/threonine kinase domain of substantial homology to CaMKs^57,58^. Genetic knockdown^57^ and chemical inhibition^59^ of these kinases have established their role in synapse maturation and dendritic outgrowth.

**Figure 4.**
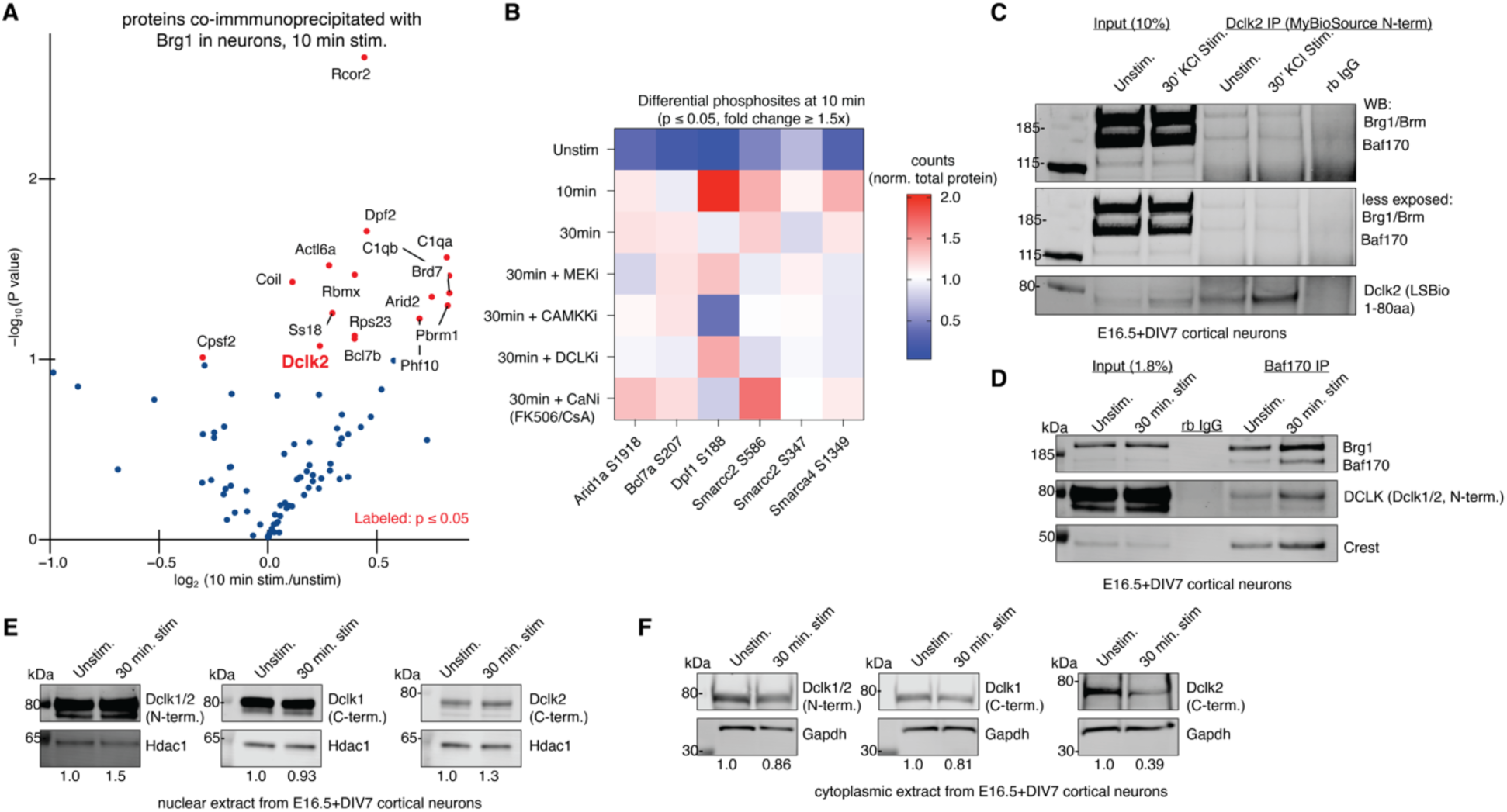
Identification of a putative BAF kinase. **A.** Differential proteins co- immunoprecipitating with Brg1 in neurons detected after 10 min. stimulation of cortical neurons; n=2 biological replicates; P-values computed by two-sided Wald test, differential sites were classified as p_adj_≤0.05. **B.** Differential phosphosites (norm. respective protein levels) detected after 10 min. stimulation of cortical neurons from Fig. 3 and their change upon addition of inhibitors of different kinases downstream of known depolarizations signaling pathways; n=2-3 biological replicates, mean of that abundance measurements are shown. **C.** Dclk2 co- immunoprecipitates Brg1 and Baf170 in cortical neurons. Two different Dclk2 antibodies used for IP and WB. Representative of n=1 biological replicates. **D.** Baf170 co-immunoprecipitates Dclk1/2 in cortical neurons. Representative of n=2 biological replicates. **E.** Changes in nuclear and **F.** cytoplasmic levels of Dclk1 and Dclk2. Representative of n=2 biological replicates. Densitometry measurements of Dclk normalized to Hdac1 or Gapdh shown below blot.

To investigate the role of Dclk in mediating BAF phosphorylation in an unbiased fashion, we treated neurons with kinase inhibitors to canonical calcium signaling pathways as well as a new selective Dclk1/2 inhibitor^59^, and assessed the consequences on activity-dependent BAF phosphorylation (Fig. 4B). Single-digit micromolar doses with short treatments (≤ 30 mins) were used to minimize off-target effects. Analysis of activity-dependent phosphosites identified in Fig. 3 revealed several that were differentially regulated by some kinase cascades compared to others. For example, Dpf1/Baf45b S188 hyper-phosphorylation was specifically reduced by CaMKK inhibition, nominating it as downstream of CaMK signaling (Fig. 4B).

Inhibition of Dclk1/2 with a pan-Dclk inhibitor^59^ substantially reduced the hyper-phosphorylation of multiple activity-dependent BAF phosphosites, most notably Baf170/Smarcc2 S586 (by almost 2-fold compared to 10 mins. depolarization) (Fig. 4B). Dclk2 co-immunoprecipitated Baf170 and Brg1 in cortical neurons (Fig. 4C). In a reciprocal co-IP from neurons with an antibody for Baf170, Dclk1/2 was robustly pulled down with a pan-Dclk antibody in two separate replicates (Fig. 4D, Supplemental Fig. 5A). These results confirmed our IP-MS finding and nominated Dclk1/2 as a potential BAF kinase that makes a stable and perhaps dedicated interaction with BAF.

### Activity-dependent nuclear translocation of Dclk

Many kinases respond to calcium signaling by shuttling to the nucleus to perform their functions^28,29^. We noticed that Dclk2 increased in our nuclear extract inputs in the co-IP experiments (Fig. 4C) and hypothesized that Dclk2 might translocate to the nucleus in a similar model of calcium-activated kinase activity. We fractionated nuclear and cytoplasmic extracts from cortical neurons before and after membrane depolarization and blotted for Dclk1/2 with three different antibodies: a pan-Dclk antibody; a Dclk1-specific antibody recognizing a C-terminal epitope, and a Dclk2-specific antibody recognizing a C-terminal epitope. Dclk2 increased ∼30% in our nuclear extracts and was depleted from the cytoplasmic extract by ∼60% (Fig. 4E,F); Dclk1, in contrast, changed only slightly (Fig. 4E,F). These experiments were repeated twice on separate biological replicates of neurons and are consistent with the observations in the nuclear extract inputs in the co-IP experiments in Fig. 4C,D. The relative abundance of Dclk2 increased by a similar degree (1.26-fold) in the nuclear extract MS (Supplemental Fig. 5B). Our data are consistent with a model where membrane depolarization activates Dclk2 translocation to the nucleus and subsequent regulation of BAF phosphorylation. We note, however, that we cannot make a statement about the role of Dclk2 in the direct phoshorylation of BAF.

## DISCUSSION

Here we show that BAF responds within minutes of neural activity by switching its composition. Relative to the core ATPase Brg1, BAF complexes switch from canonical BAF to PBAF, containing Arid2/Baf200, Pbrm1/Baf180, Brd7, and Phf10/Baf45a. The switch occurs in two different cell types (fibroblasts and neurons) and is downstream of four different canonical growth factor and calcium signaling pathways.

DNA sequence motifs associated with nBAF complexes after neural activity were significantly (q < 0.0001) enriched for X-box/RFX TF binding motifs within 10 minutes of depolarization. A biochemical interaction between BAF and RFX TFs could not be detected, but PBAF uniquely contains an RFX DNA-binding domain. We strongly hypothesize that the compositional switch from BAF to PBAF governs BAF localization and remodeling function; however, further analyses of PBAF binding on chromatin should be performed to test this theory.

We also identify a selective set of 6 activity-dependent hyper-phosphorylations on BAF complex subunits, that change up to 20-fold within 10 minutes of membrane depolarization. A functional analysis of one such phosphosite, on Baf170/Smarcc2, suggested that constitutive phosphorylation of it directed Sox TF binding activity and neural development. We identify a novel BAF-interacting, neural-specific kinase, Dclk2, whose inhibition reduces the phosphorylation of multiple activity-dependent BAF phosphosites, including on Baf170. We found that Dclk2 also translocates into the nucleus upon membrane depolarization, suggesting a regulatory role in controlling BAF activity and or composition. However, direct demonstration of the activity of Dclk2 will require future work with purified proteins and mutations in both Dclk2 and its putative sites in BAF.

In summary we provide the first evidence that BAF complexes respond molecularly to neural activity within minutes, similar to other activity-responsive TFs such as CREB, SRF, and MEF2. By mapping the consequences on chromatin of activity-dependent effects on BAF, we also outline how a chromatin remodeler can respond to membrane signaling to act immediately on chromatin. We hope these studies will assist investigations into the molecular mechanisms by which BAF mutations cause neurodevelopmental disorders.

## LIMITATIONS OF THE STUDY

This work models neuronal activity in cultures of embryonic primary mouse cortical neurons, which are removed from their tissue environments containing other cell types and purified. Future work could explore the molecular consequences of activity in an animal, which would contain other brain cell types that might affect activity-dependent responses. In addition, in this study activity was modeled using membrane depolarization by 50 mM KCl, an established protocol but one whose limitations have been noted^60^. KCl-mediated depolarization also activates many different intracellular signaling cascades. Future work could explore the link between the findings on BAF regulation reported here and specific signaling pathways, using, for example, optogenetic approaches to stimulate specific categories of neurons or the addition of particular neurotransmitters.

## METHODS

### Animal Husbandry

In all animal experiments, mice used were wild-type CD-1 outbred mice, housed at Stanford up to 5/cage in a colony with a 12-hour light/dark cycle (lights from 7:00 am to 7:00 pm) at constant temperature (23 °C) with ad libitum access to water and food. Animal protocols were approved by IACUC at Stanford University under protocol number 8801.

### Primary neuron culture

Embryonic cortical neurons were harvested from E16.5 mice. Pregnant female mice were anesthetized with isoflurane and euthanized by cervical dislocation on E16.5. Cortical neurons were cultured following a protocol described in^9^. Briefly, culture plates (Corning #353003) were coated overnight at 37 °C with poly-D-lysine (Sigma #P6407) at 0.1 mg/mL in borate buffer (1.55 g boric acid/2.4 g sodium tetraborate in 500 mL sterile water). Plates were washed 3x with water and 1x with PBS before plating neurons. Embryonic cortices were harvested by microdissection and enzymatically dissociated to single neurons with Neural Tissue Dissociation Kit (P) (Miltenyi Biotech #130-092-628). Cortical neurons from separate pups were kept distinct as biological replicates. Neurons were resuspended in plating media [DMEM (Life Technologies #11960-077), 10% FBS (Omega #FB-01), 1x penicillin-streptomycin (ThermoFisher #15140122)], counted, and plated at a density of 30M neurons per 15 cm plate, 10M per 10 cm plate, 1M per well of a 6-well plate, 0.5M per well of a 12-well plate, or 0.1M per well of a 24-well plate. After incubation for 1 hour at 37 °C, neurons were attached to the plates and media was exchanged with neuron maintenance media [Neurobasal Medium (Life Technologies #21103049), 2.5% B27 (ThermoFisher #17504044), 1% penicillin-streptomycin (Life Technologies #15070-063), 0.25% GlutaMax (Life Technologies #35050061)]. On day in vitro 2 (DIV2), ara-c (Sigma #C1768) was added to a final concentration of 0.5 µM to inhibit proliferation of any contaminating glial cells. Beginning on DIV3, half the media was exchanged for fresh neuron maintenance media every three days.

### Membrane Depolarization by KCl Stimulation

At the designated DIV, neurons were silenced overnight with 1 µM tetrodotoxin (Tocris #1069) and 100 µM D-AP5 (Tocris #79055-68-8) to reflect the resting state^61^ and then depolarized with 50 mM KCl for indicated time points. Neurons were then immediately moved to ice, media was then immediately exchanged to ice-cold PBS, and harvested for downstream assays.

### Mouse Embryonic Fibroblast (MEF) cell culture

Mouse embryonic fibroblasts were generated from F2 pups at E16.5 according to established procedure in^62^ and immortalized with SV40 T antigen. MEFs were cultured in DMEM + 10% FBS + 1x penicillin-streptomycin. Cultures were passaged by trypsinization and were used before passage 10. Cultures were routinely checked for mycoplasma and immediately checked upon suspicion. No cells tested positive.

### MEF Serum Stimulation

MEFs were serum starved in 0.5% FBS for 24 hours before stimulation with serum up to a final concentration of 15% for indicated time points. Stimulation was stopped by immediate movement of the cultures to ice, washing with ice-cold PBS plus 10mM sodium butyrate to inhibit histone deacetylases, 1mM sodium orthovanadate, and 1:1000 v/v protease inhibitor cocktail (chymostatin, Millipore #230790; leupeptin, Millipore #108975; pepstatin, Millipore #516481) and 1 phosphatase inhibitor tablet/10 mL (PhosSTOP, Roche #4906837001), and harvesting by scraping with a cell lifter.

### CRISPR constructs

CRISPR knockout constructs were constructed using lentiCRISPR v2 (a gift from Feng Zhang, Addgene plasmid # 52961; http://n2t.net/addgene:52961; RRID:Addgene_52961) using gRNAs targeting isoform-conserved exons in mouse *Smarcc2/*Baf170. 3 gRNAs were designed using CRISPick^63^ and pooled virus was made from the 3 lentiCRISPR v2 plasmids. Target sequences were: (1) GCGTCCATGCCATTGAACGG; (2) ACACCGACACATTCAACGAG; and (3) GACAGGATACACAACATGGG.

### Overexpression constructs

Baf170 overexpression constructs were subcloned from Baf170 cDNA published previously in^53^, into an EF1a-driven lentiviral vector. Briefly, first, the Baf170 wild-type overexpression construct was cloned by wobbling bases corresponding to the PAM sequences targeted by the Baf170 gRNAs, using site-directed mutagenesis (NEB #E0554S). Then, further site-directed mutagenesis was performed to generate point mutants at phosphosites. All constructs were verified using Sanger sequencing.

### Lentivirus production

Lentivirus was produced using a standard protocol^64^ from a 15 cm plate of 25M lenti-x 293T cells (Clontech #632180) via polyethylenimine (PEI) transfection, using lentiviral constructs and packaging plasmids psPAX2 and pMD2.G. Two days after transfection, the media containing the virus was collected, filtered with a 0.4 µM filter, and ultra-centrifuged for 2 hours at 20,000 rpm in an SW28 rotor (Beckman). Viral pellets were resuspended in 250 µL PBS and flash-frozen. Relative titer was estimated by Lenti-X GoStix Plus (Takara #631280) and was similar for all viral constructs.

### Transduction of neurons

Virus was applied to wells containing primary neurons overnight, and media was exchanged for virus-free media the next day. A total of 20 µL of virus was applied to one well of a 12-well plate of 0.5M neurons/well, and 50 µL was applied to one well of a 6-well plate of 1M neurons/well.

For knockout and overexpression of Baf170 mutants, virus aliquots for both the knockout constructs and the overexpression constructs were applied at the same time in equal volumetric ratios. Virus was added in DIV4 and cells harvested on DIV18 to allow time for overexpression and knockout, based on past experience of how long it took for effective transduction in neurons.

### Kinase Inhibition Strategy

Inhibitors were chosen to inhibit canonical calcium-activated kinase signaling pathways at their kinases closest to the nucleus that have been identified, and have specific, cell-permeable, small molecule inhibitors. Compounds and concentrations used were: CaMKKi: STO-609, 3 µM; MEK1/2i: PD0325901, 3 µM; DCLKi: DCLK-IN-1, 2.5 µM; CaNi: 10 nM FK506 + 1 µM cyclosporin (CsA). Cyclosporin A and FK506 were used together because of the greater abundance of calcineurin in neurons compared to FKBP and cyclophilin, which are required for formation of inhibitor complexes^30^. Inhibitors were given at a final concentration of 0.1% DMSO for a 30 minute pre-treatment before depolarization or rest for the indicated times (30 min or less).

### Nuclear Extract Preparation from Neurons

A 15 cm plate of neurons was washed 1x with 10 mL ice cold PBS plus 10mM sodium butyrate to inhibit histone deacetylases, 1mM sodium orthovanadate, and 1:1000 v/v protease inhibitor cocktail (chymostatin, Millipore #230790; leupeptin, Millipore #108975; pepstatin, Millipore #516481) and 1 phosphatase inhibitor tablet/10 mL (PhosSTOP, Roche #4906837001). Cells were lifted with a cell lifter and centrifuged for 2000 g for 5 min at 4°C. Pellets were flash-frozen and stored until further processing. Then, after thawing on ice, cells were washed once more with 10 mL cold PBS plus 10mM sodium butyrate to inhibit histone deacetylases, 1mM sodium orthovanadate, and protease and phosphatase inhibitors (hereafter: PBS*) and resuspended in 250 µL Buffer A (25mM HEPES pH 7.5, 25mM KCl, 5mM MgCl_2_, 10% glycerol, 0.1% NP-40) and 10mM sodium butyrate to inhibit histone deacetylases, 1mM sodium orthovanadate, and protease and phosphatase inhibitors (hereafter: Buffer A*), incubated for 7 mins on ice, and an aliquot taken to count nuclei using Trypan Blue and a hematocytometer. Nuclei were spun down at 1500 rpm for 5 min at 4°C, and the supernatant saved as the cytosolic fraction. Nuclei were then resuspended in 5 mL Buffer A*, incubated on ice for 7 minutes again, and another aliquot was checked by a hematocytometer to make sure >95% were nuclei. Nuclei were then spun down again at 1500 rpm for 5 min at 4°C, washed 1x with 5 mL Buffer C (10mM HEPES pH 7.5, 100mM KCl, 2mM MgCl_2_, 10% glycerol, 0.5mM CaCl_2_) + 10mM sodium butyrate to inhibit histone deacetylases, 1mM sodium orthovanadate, and protease and phosphatase inhibitors (hereafter: Buffer C*). Nuclei were finally resuspended in 700 µL Buffer C* and chromatin was precipitated out by slow dropwise addition of 108 µL 3M ammonium sulfate and rotation at 4°C for 1 hour. Chromatin was spun down by ultracentriguation in polyallomer centrifuge tubes (Beckman #343778 11x34mm) at 100,000 rpm for 15 mins at 4°C and the supernatant was saved. The concentration of protein in the supernatant was measured by Bradford. For mass spectrometry analyses, the nuclear extract was precipitated by 1:4 v/v addition of trichloroacetic acid (TCA), vortexing and incubation on ice for 30 mins, pelleting by centrifugation at 16,000 g for 15 mins at 4°C, washing 1x with 1 mL ice-cold acetone, and finally washing 2x with 1 mL ice-cold methanol. Pellets were stored at -80°C until ready for injection. For SDS-PAGE analyses, nuclear extracts were equalized in protein concentration and mixed with 4X NuPage LDS loading buffer (Life Technologies, cat# NP0007) and 1mM DTT.

### Ammonium sulfate precipitation of soluble nuclear protein

After nuclear extraction and chromatin precipitation with ammonium sulfate as above, 233 mg of solid ammonium sulfate was added to the 805 µL of supernatant (in Buffer C*) and resuspended well, then incubated on ice for 40 minutes with mixing every 10 minutes. BAF complexes and other nuclear proteins were precipitated by ultracentrifugation at 100,000 rpm for 15 mins at 4°C and the pellet was saved at -80°C until immunoprecipitation.

### BAF Immunoprecipitation (IP) from Neurons

Precipitated nuclear protein was resuspended in 205 µL IP Buffer (20mM HEPES pH 7.5, 150 mM KCl, 1mM MgCl_2_, 0.5mM CaCl_2_, 10% glycerol, 0.1% Triton X-100) + 10mM sodium butyrate to inhibit histone deacetylases, 1mM sodium orthovanadate, and protease and phosphatase inhibitors (hereafter: IP Buffer*) and concentration was measured by Bradford. 8 µg was saved for input and 256 µg was used per IP at 0.86 µg/µL, with overnight rotation at 4°C. For each BAF IP, 5 µg SantaCruz H-10 mouse monoclonal Brg1 antibody was used, crosslinked to Protein G Dynabeads (Invitrogen #10003D) using BS3 (bis(sulfosuccinimidyl)suberate) (ThermoFisher #A39266): 5µg antibody was incubated with rotation at room temperature to 50 µL Protein G beads, washed 2X with 1 mL PBS and 2X with 1 mL Conjugation Buffer (20mM sodium phosphate, 150mM NaCl pH 8.0), then incubated with 230 uL of 2.86 mg/mL BS3/Conjugation Buffer for 30 mins exactly at RT under rotation, quenched with 12.5 µL 1M Tris pH 7.5 with 15 mins rotation at RT, and washed 2X with IP*. IPs were washed 3X the next day with 1mL IP Wash* (IP* except 300mM KCl instead of 150mM KCl) and eluted with a low-pH/glycine protocol: beads were resuspended in 50 µL 0.1M glycine-HCl pH 2.5, incubated with shaking at 900 rpm for 30 mins at 37°C, centrifuged briefly then supernatant was transferred to a new tube on ice and neutralized with 5 µL 1M Tris pH 8, and repeated 1X. 10% was run on a gel to confirm the IP and 90% was TCA-precipitated as above for mass spectrometry analyses.

### Co-immunoprecipitation of BAF and Dclk

Immunoprecipitation was carried out as above except 5 µg Baf170 antibody (homemade rabbit polyclonal raised against conserved epitope around Ile818) or Dclk2 (MyBioSource MBS9604810, recognizes N-terminal region) was used, non-crosslinked. In both cases IP was cross-validated using immunoblot against the same bait but with a different antibody: Baf170 (SantaCruz monoclonal mouse E-6) or Dclk2 (LSBio LS-C185753, recognizes N-terminal region), respectively.

### Glycerol Gradient

Glycerol gradients (10-30%) of nuclear proteins were carried out as previously established^65^. Briefly, neuronal or MEF nuclear extract was resuspended in ice-cold HEMG-0 buffer (50 mM HEPES pH 7.9, 100mM KCl, 0.1 mM EDTA, 12.5 mM MgCl_2_) + 1mM sodium orthovanadate, 10mM sodium butyrate, and protease and phosphatase inhibitors (hearafter: HEMG-0*). 2.5% was reserved as input. A 10-30% gradient of glycerol in HEMG buffer was poured into 14 x 89 mm polyallomer centrifuge tubes (Beckman #331372). The resuspended nuclear extract was carefully layered on top of the gradient and centrifuged in an SW41 rotor at 40,000 rpm for 16 hours at 4°C. Twenty 500 µL fractions were collected carefully, without disturbing the gradient, and each fraction was TCA/acetone precipitated, by 1:4 v/v addition of trichloroacetic acid (TCA), vortexing and incubation on ice for 30 mins, pelleting by centrifugation at 16,000 g for 15 mins at 4°C, washing 1x with 1 mL ice-cold acetone, and finally washing 2x with 1 mL ice-cold methanol. Fractions were then resuspended in equal volumes of LDS loading buffer (Life Technologies, cat# NP0007) and 1mM DTT and run on SDS-PAGE gels. Proteins were transferred at constant 85 mA, 4°C overnight to PVDF membranes and blotted using antibodies to subunits of the BAF complex or other controls.

### Sample processing for quantitative TMT-proteomic analysis

Samples were prepared following the SL-TMT protocol^66^. A total of 100 µg of protein from each nuclear extract sample (for total nuclear extract MS proteomics) or the elute from the Brg1 immunoprecipitation (for the BAF IP-MS) was used. Reduction of sample (5 mM TCEP for 15 min.) was followed by alkylation (10 mM iodoacetamide for 30 min.) and quenching (5 mM DTT for 15 min.). Samples were then chloroform-methanol precipitated. The precipitated proteins were resuspended in 200 mM EPPS pH 8.5, digested first by Lys-C overnight at room temperature and later by trypsin (6 h at 37°C). Both enzymes were used at a 1:100 enzyme-to-protein ratio.

The samples were then labeled with tandem mass tag (TMTpro) reagents^66^. Acetonitrile was added to a final volume of 30% prior to adding the TMTpro labeling reagent. For protein level analysis, ∼50 µg of peptides were labeled with 100 µg of TMT. For phosphopeptide analysis, we estimated the phosphopeptide enrichment to be ∼1.5:100 and so ∼30 µg of peptides were labeled with 60 µg of TMT. Labeling occurred at room temperature for 1 h. Once labeling efficiency was verified (here, >97%), hydroxylamine was added at a final concentration of ∼0.3% and incubated for 15 min. at room temperature and the samples were pooled at a 1:1 ratio across all channels.

To enrich phosphopeptides, the pooled sample was desalted over a 200 mg SepPak column and phosphopeptides were enriched with the Pierce High-Select Fe-NTA Phosphopeptide enrichment kit following manufacturer’s instructions. The eluate was desalted via StageTip^67^ and was ready for MS analysis. The washes and the unbound fraction of this enrichment were desalted and used for proteome-level analysis.

For total nuclear extracts, the pooled sample was fractionated using basic-pH reversed-phase (BPRP) Liquid Chromatography using an Agilent 1200 pump with an Agilent 300Extend C18 column (2.1 mm ID, 3.5 μm particles, and 250 mm in length). The flow rate over the column was 0.25 mL/min and we used 50-min linear gradient with 5% to 35% acetonitrile in 10 mM ammonium bicarbonate pH 8. Ninety-six fractions were collected and concatenated into 24 superfractions prior to desalting^68^. These 24 superfractions were sub-divided into two groups, each consisting of 12 non-adjacent superfractions. These superfractions were subsequently acidified with 1% formic acid and vacuum centrifuged to near dryness. Each superfraction was desalted via StageTip^67^. Once dried by vacuum centrifugation, the sample was reconstituted using 5% formic acid and 5% acetonitrile prior to acquisition of LC-MS/MS data.

### Mass Spectrometry Data Collection and Processing

Mass spectrometric data were collected on an Orbitrap Fusion Lumos mass spectrometer, both which are coupled to a Proxeon NanoLC-1200 UHPLC and a FAIMSpro interface^69^. The 100 μm capillary column was packed with 35 cm of Accucore 150 resin (2.6 μm, 150 Å; ThermoFisher Scientific).

Mass spectrometric data for total nuclear extracts were collected on Orbitrap Fusion Lumos instruments (using RTS-MS3) coupled to a Proxeon NanoLC-1200 UHPLC. The 100 µm capillary column was packed with 35 cm of Accucore 150 resin (2.6 μm, 150Å; ThermoFisher Scientific) at a flow rate of 565 nL/min. The scan sequence began with an MS1 spectrum (Orbitrap analysis, resolution 60,000, 400−1600 Th, automatic gain control (AGC) target 100%, maximum injection time “auto”). Data were acquired ∼90 minutes per fraction. MS2 analysis consisted of collision-induced dissociation (CID), quadrupole ion trap analysis, automatic gain control (AGC) 100%, NCE (normalized collision energy) 35, q-value 0.25, maximum injection time 35ms), and isolation window at 0.7 Th. RTS was enabled and quantitative SPS-MS3 scans (resolution of 50,000; AGC target 2.5x105; collision energy HCD at 55%, max injection time of 250 ms) were processed through Orbiter with a real-time false discovery rate filter implementing a modified linear discriminant analysis. For FAIMS, the dispersion voltage (DV) was set at 5000V, the compensation voltages (CVs) used were -40V, -60V, and -80V, and -30V, -50V, and - 70V, and the TopSpeed parameter was set at 1 sec.

For phosphopeptide profiling, data were acquired using three to five injections on an Orbitrap Eclipse mass spectrometer with varying combinations of FAIMS CVs between -30 and -80V (3 CVs per set) over a 150 min gradient. A 1sec TopSpeed cycle was used for each CV. The scan sequence began with an Orbitrap MS1 spectrum with the following parameters: resolution: 120,000, scan range: 400-1500 Th, automatic gain control (AGC): 100%, and maximum injection time: “auto.” MS2 analysis consisted of higher-energy collisional dissociation (HCD) with the following parameters: resolution: 50,000, AGC: 300%, normalized collision energy (NCE): 36%, maximum injection time: 250 ms, and isolation window: 0.5 Th, and. In addition, unassigned, singly, and > 5+ charged species were excluded from MS2 analysis and dynamic exclusion was set to 60 s.

For IP-MS profiling, data were acquired using multiple injections (n=6) on an Orbitrap Eclipse mass spectrometer with varying combinations of FAIMS CVs between -30 and -80V (3 CVs per set) over a 150 min gradient. A 1sec TopSpeed cycle was used for each CV. The scan sequence began with an MS1 spectrum (Orbitrap analysis, resolution 60,000, 400-1500 Th, automatic gain control (AGC) target 100%, maximum injection time “auto”). Data were acquired ∼90 min per fraction. The hrMS2 stage consisted of fragmentation by higher energy collisional dissociation (HCD, normalized collision energy 36%) and analysis using the Orbitrap (AGC 300%, maximum injection time 200 ms, isolation window 0.5 Th, resolution 50,000).

Once the spectra were converted to mzXML using MSconvert^70^, database searching could be performed. Database searching included all mouse entries from UniProt (downloaded March 2021). which was concatenated with a version of the database in which the order of the amino acid residues of each protein was reversed. Database searches used a 50-ppm precursor ion tolerance and a product ion tolerance of 0.03 Da for hrMS data. RTS-MS3 data used a 0.9 Da product ion tolerance^71,72^. For static modifications, lysine residues and peptide N-termini were modified with +304.207 Da due to the TMTpro labels and +229.162 Da for classic TMT (IP-MS only). Meanwhile, all cysteine residues were modified with iodoacetamide (carbamidomethylation) that results in a +57.021 Da increase in mass. Also, methionine oxidation (+15.995 Da) was set as a variable modification. Likewise, deamidation (+0.984 Da) at glutamine and asparagine residues and phosphorylation (+79.966 Da) at serine, threonine, and tyrosine residues were also set as variable modifications for phosphopeptide enrichment. The false discovery rate (FDR) was set at 1% at the peptide level with filtering a linear discriminant analysis^73^. The protein lists were assembled further to a final protein-level FDR of 1%. The intensities of reporter ions were corrected for the isotopic impurities of the different TMT reagents^74^. For each protein, the peptide summed signal-to-noise (S/N) measurements were summed and normalized to account for equal protein loading by equating the sum of the signal for all proteins in each channel. For BAF IP-MS, this was normalized by the relative abundance for the bait, Brg1. For phosphosite identification, the AScore^71^ false-discovery metric was used and only phosphosites that were “high-confidence”, with p ≤ 0.05, were retained. Students’ t-test was used for differential site and protein pull-down calling. Human keratin and other contaminants were excluded from downstream analyses (contaminants made up <5% of all peptides).

### RNA Extraction, qPCR, and Sequencing Library Preparation

Cells were harvested in TRIsure (Bioline #38033). RNA was extracted using Direct-zol RNA MicroPrep columns (Zymo #R2062) treated with DNAseI. cDNA was prepared for RT-qPCR using the SensiFAST cDNA preparation kit according to manufacturer instructions (Bioline #65054). 1µL of cDNA was used per RT-qPCR reaction prepared with SYBR Lo-ROX (Bioline #94020). *cFos* primers were from^9^: *Fos* FWD: CGGGTTTCAACGCCGACTA *Fos* REV: TTGGCACTAGAGACGGACAGA; *Gapdh* FWD: CTGACGTGCCGCCTGGAGAAAC *Gapdh* REV: CCCGGCATCGAAGGTGGAAGAGT. For sequencing library preparation, rRNA was depleted using (NEB #6350) and prepared into paired-end libraries (NEB #E7760S). Libraries were sequenced on an Illumina NovaSeq (Novogene).

### ATAC-seq

ATAC-seq libraries were constructed from on-plate nuclear prep of neurons plated at 1M/6-well of a 6-well plate. Briefly, DNase at a final concentration of 200 U/mL was added to wells for 30 mins at 37°C. After indicated treatment timepoints (such as stimulation, and/or drug addition [100 nM FHT1015 or DMSO at final 0.1% DMSO concentration]), neuronal media was quickly dumped and neurons were washed 4X with cold PBS. 1mL of RSB-Lysis buffer/well (10mM Tris pH 7.4, 10mM NaCl, 3 mM MgCl_2_, 0.1% NP-40, 0.1% Tween-20, 0.01% Digitonin) was added and incubated for 10 mins on ice, then nuclei were gently lifted, counted by a hematocytometer, and spun down at 500 g for 4°C for 10 mins in a swing-bucket centrifuge. 75,000 nuclei were taken for further processing. Lysis buffer was washed out with RSB-Wash (containing 0.1% Tween-20 but no NP-40 or Digitonin). The pellet was resuspended in transposition mixture (25 µL 2X TD Buffer (20mM Tris pH 7.6, 10mM MgCl_2_, 20% DMF), 100 nM final Tn5 transposase (homemade, gift from William Greenleaf), 16.5 µL PBS, 0.5 µL 1% digitonin, 0.5 µL 10% Tween- 20, 5 µL H_2_O) and incubated for 30 mins at 37°C with 1000 rpm mixing. Reactions were cleaned up, amplified into libraries, and quantified using published protocol^75^. Libraries were sequenced on an Illumina NovaSeq (Novogene).

### ATAC-seq Analysis

ATAC-seq reads were checked for quality using fastqc (https://www.bioinformatics.babraham.ac.uk/projects/fastqc/) and trimmed from adapters using trim_galore (https://github.com/FelixKrueger/TrimGalore) with parameters --paired --nextera. Trimmed reads were aligned to the mouse mm10 genome using bowtie2 with parameters -- very-sensitive -X2000. Low quality reads, duplicated reads and reads with multiple alignments were removed using samtools^76^ and Picard (https://broadinstitute.github.io/picard/). Positions of Tn5 inserts were determined as read start position offset by +4 bp for reads aligned to the + strand and as a read start position offset by -5 bp for reads aligned to the - strand. macs2^77^ callpeak was used for peak calling with Tn5 insert positions and parameters -p 0.01 –nolambda --shift -75 --extsize 150 --nomodel --call-summits --keep-dup all. Bedtools^78^ was used to find consensus set of peaks by merging peaks across multiple conditions (bedtools merge), count number of reads in peaks (bedtools intersect -c) and generate genome coverage (bedtools genomecov -bga). The peak differential analysis and PCA analysis was performed using DESeq2^79^. Motif enrichment analysis was performed on peak summits using findMotifsGenome.pl from HOMER^80^ using parameters mm10 -size 200 -bg $bgfile where $bgfile = background of all consensus peaks detected. In a parallel pipeline to check the reproducibility of the analyses, bam files generated after bowtie2 alignment were processed using ChrAccR suite with default parameters, which uses 200 bp tiling windows to summarize the count data, (https://greenleaflab.github.io/ChrAccR/articles/overview.html), to identify footprints, and motifs that drive variability using ChromVAR^36^.

### RNA-seq Analysis

Raw reads were checked for quality using fastqc (https://www.bioinformatics.babraham.ac.uk/projects/fastqc/) and trimmed from adapters using cutadapt^81^ using parameters cutadapt -a AGATCGGAAGAGCACACGTCTGAACTCCAGTCA -b AGATCGGAAGAGCGTCGTGTAGGGAAAGAGTGT --nextseq-trim=20 --minimum-length 1. Transcripts were quantified using kallisto^82^. Differential gene analysis was performed using DESeq2^79^ using apeglm^83^ to shrink log_2_FoldChanges and pathway and enrichment analyses using Enrichr^84^.

### Activity-dependent Dendritic Outgrowth Experiment and Analysis

Activity-dependent dendritic outgrowth was studied using established protocol in^6^. Briefly, after dissection, cortical neurons were nucleofected (Lonza #VPG-1001) with knockout, vector control, or overexexpression constructs, along with an IRES-puro-GFP construct^53^ to enable visualization, and plated at 50,000/well on poly-D-lysine-coated 24-well glass plates (Cellvis #P24-1.5H-N). The same amount of DNA/cells were added to each mutation. On DIV4, wells were stimulated with 30 mM KCl final overnight (18 hours) or silenced with D-AP5/TTX. The next day, neurons were washed 1X with PBS by incubation for 5 mins at RT, fixed in 4% PFA, washed 1X in PBS, permeabilized with 0.3% Triton-X-100/PBS for 15 mins at RT, washed 3X with PBS, blocked with 2.5% normal donkey serum (Jackson ImmunoResarch #005-000- 121)/2.5% normal goat serum (Jackson ImmunoResearch #017-000-121)/1% BSA for 1 hour at RT, then stained with 1:2000 chicken anti-GFP (Aves #GFP-1020) for visualization of dendrites and 1:3200 rabbit anti-cFos (Cell Signaling 9FG) to validate stimulation, overnight at 4°C. After 3X PBS wash, respective secondary antibodies were added at 1:1000, washed 3X in PBS, and imaged using Keyence BZ-X710 at 40X magnification. Dendrites were manually traced from the center of the soma using SNT^85^, with 20-30 neurons collected per replicate where the analyzer was blinded to the condition being analyzed (labeled only by a code by another experimenter). Statistics including Sholl analysis and branch lengths were collected using SNT’s Sholl Analysis command and Batch→Measure Multiple Files command. Statistics were plotted using GraphPad Prism.

### Western Blotting

Cells were harvested on ice in RIPA buffer (50mM Tris-HCl pH 8, 150mM NaCl, 1% NP-40, 0.1% DOC, 1% SDS, protease inhibitor cocktail (homemade), 1mM DTT) and 1:200 benzonase (Sigma #E1014) was added and incubated for 20 mins. After 10 min centrifugation at 14,000g and 4 °C, the supernatant was collected and protein concentration was measured by Bradford. Antibodies used for immunoblots are: Brg1 (Santa Cruz H-10), Brg1/Brm (homemade rabbit polyclonal, J1), Baf170 (homemade rabbit polyclonal, raised against conserved epiptope around Ile818), Baf170 (SantaCruz mouse monoclonal E-6), Vinculin (ThermoFisher 700062), DCLK (homemade rabbit polyclonal, raised against conserved N-terminus present in both Dclk1 and Dclk2, kind gift from Prof. Joseph Gleeson’s laboratory), Dclk1 (Cell Signaling 14361, recognizes C-terminus of Dclk1), Dclk2 (LSBio LS-C185753, recognizes N-terminal region), Dclk2 (MyBioSource MBS9604810, recognizes N-terminal region), Dclk2 (abcam 106639, recognizes C-terminal region), CREST (SantaCruz D-7), HDAC1 (Cell Signaling 10E2), and GAPDH (Santa Cruz 6C5). All antibodies were used at 1:1000 v/v dilutions except GAPDH (1:2000). ImageStudio (Licor) was used for blot imaging.

## ACKNOWLEDGEMENTS

S.G. would like to thank L. Chen for advice in neuronal dissections and mouse husbandry, C.Weber for advice in organizing the proteomic analyses, N.S.G. for providing the Dclk inhibitor, and E. Raz and A. Krokhotin for advice in performing the transcription factor motif analyses. All authors would like to thank members of the Crabtree lab for input and advice. The studies described in this manuscript were funded from a grant from HHMI to G.R.C., NIH grants CA276167, CA163915 and MH126720-01 to G.R.C. Funding was provided to S.G. from NIH grant 5F31HD103339-03. This work was funded in part by NIH/NIGMS grants R01 GM67945 to S.P.G. and R01 GM132129 to J.A.P.

## AUTHOR CONTRIBUTIONS

S.G., W.W., and G.R.C. conceived the project. S.G. performed neuronal dissections, molecular biology, biochemical, and genomics experimental and computational analyses. W.W. assisted with neuronal dissections, biochemical experiments, and project design. J.P. and S.P.G. performed mass spectrometry (phospho)proteomics with extracts provided by S.G. C.L. assisted with cloning under supervision of S.G. S.H.K. performed the Baf170 mutant ATAC-sequencing and the cortical neuronal RNA-sequencing together with S.G. K.R. analyzed activity-dependent dendritic outgrowth under supervision of S.G. S.G. and G.R.C. wrote the manuscript with input of all authors.

## DATA AVAILABILITY

Sequencing data is deposited to GSE245256. The mass spectrometry proteomics data have been deposited to the ProteomeXchange Consortium via the PRIDE^86^ partner repository with the dataset identifier PXD046031. All other materials and reagents available from authors upon request.

## SUPPLEMENTARY INFORMATION

**Supplemental Figure 1.**
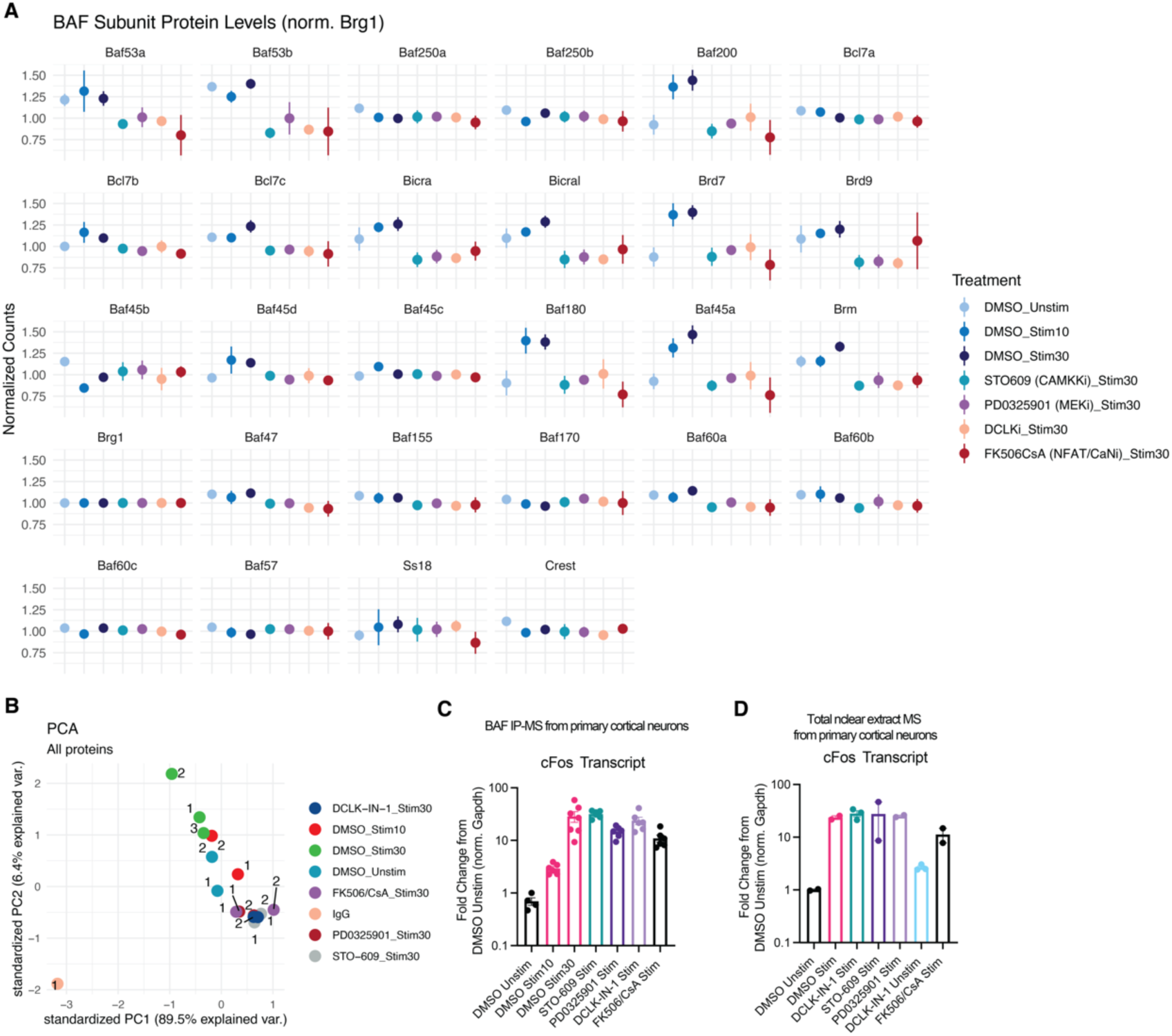
Proteomic characterization of BAF complexes in cortical neurons. **A.** All BAF subunits immunoprecipitated with Brg1 as a function of time and drug; mean±s.d., n=2-3 biological replicates, abundance normalized to Brg1 abundance. **B.** Principal component analysis of all proteins co-immunoprecipitated with Brg1; n=2-3 biological replicates, abundance normalized to Brg1 abundance. **C.** Validation of stimulation protocol for IP-MS by measurement of *cFos* mRNA by RT-qPCR; mean±s.d., n=2-3 biological replicates with 2-3 technical replicates each. **D.** Validation of stimulation protocol for whole nuclear extract MS by measurement of *cFos* mRNA by RT-qPCR; mean±s.d., n=2-3 biological replicates with 2-3 technical replicates each.

**Supplemental Figure 2.**
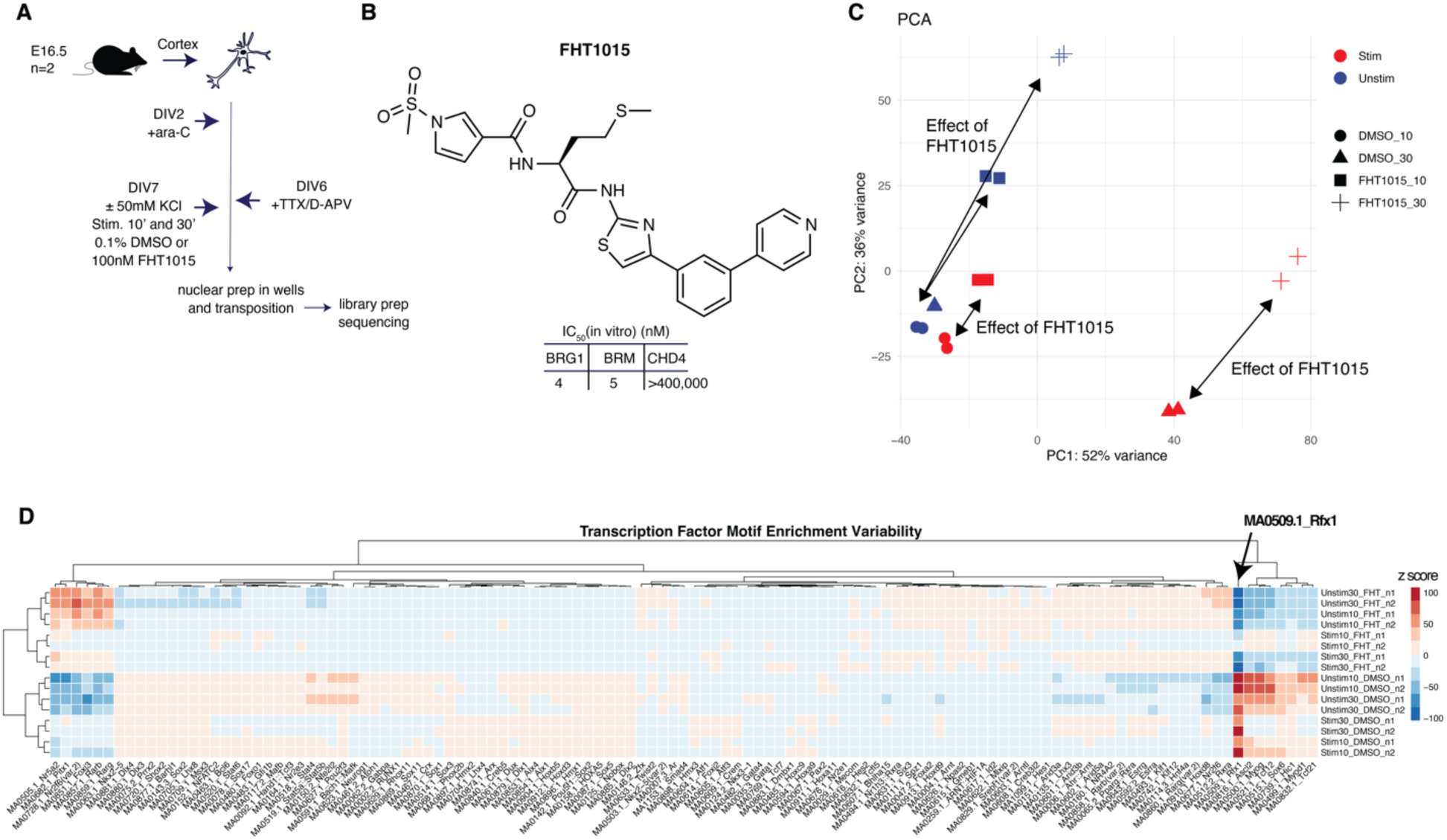
Activity-dependent chromatin accessibility changes regulated by BAF complexes. **A.** Schematic of ATAC-seq experiment. **B.** Structure of BAF ATPase inhibitor FHT1015 and reported IC_50_ values for ATPases. **C.** Principal component analysis of ATAC-seq experiment in **A. D.** Analysis of transcription factor motif variability across all conditions by ChromVAR^36^.

**Supplemental Figure 3.**
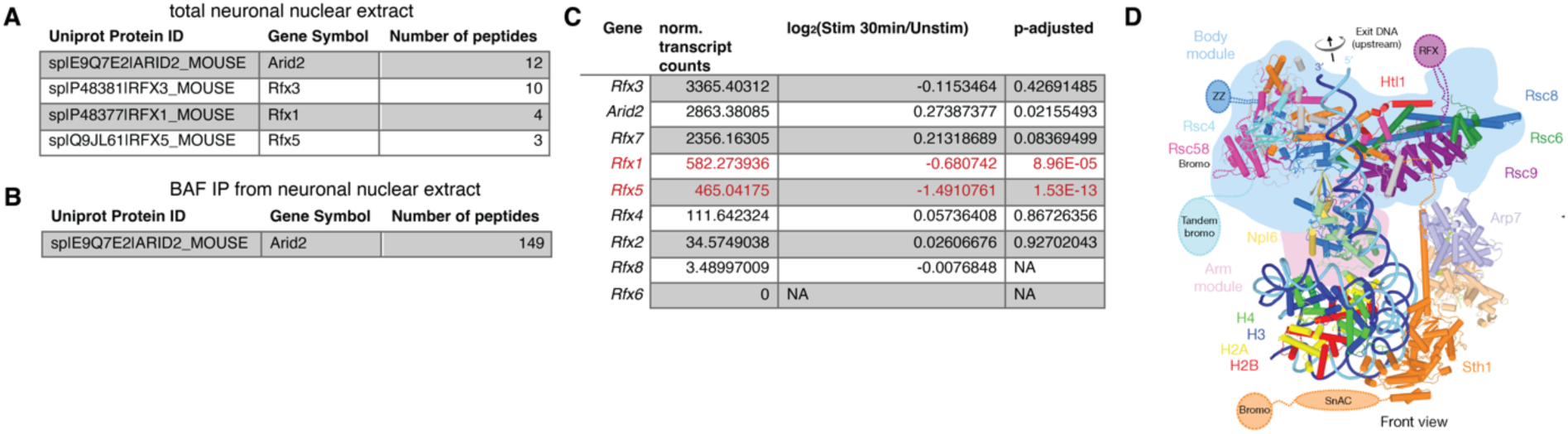
Characterization of RFX transcription factors in neurons. **A.** Number of peptides of all Rfx-family transcription factors and Arid2 detected in total nuclear extract from cortical neurons. **B.** Number of peptides of all Rfx-family transcription factors and Arid2 detected in all Brg1-immunoprecipitated proteins from cortical neurons. **C.** Normalized, transcript-isoform-collapsed abundance measurements of *Rfx* genes in cortical neurons and changes observed after stimulation. **D.** Cryo-EM structure of RSC, the yeast homolog of PBAF, displaying density for the RFX domain near extra-nucleosomal DNA^48^.

**Supplemental Figure 4.**
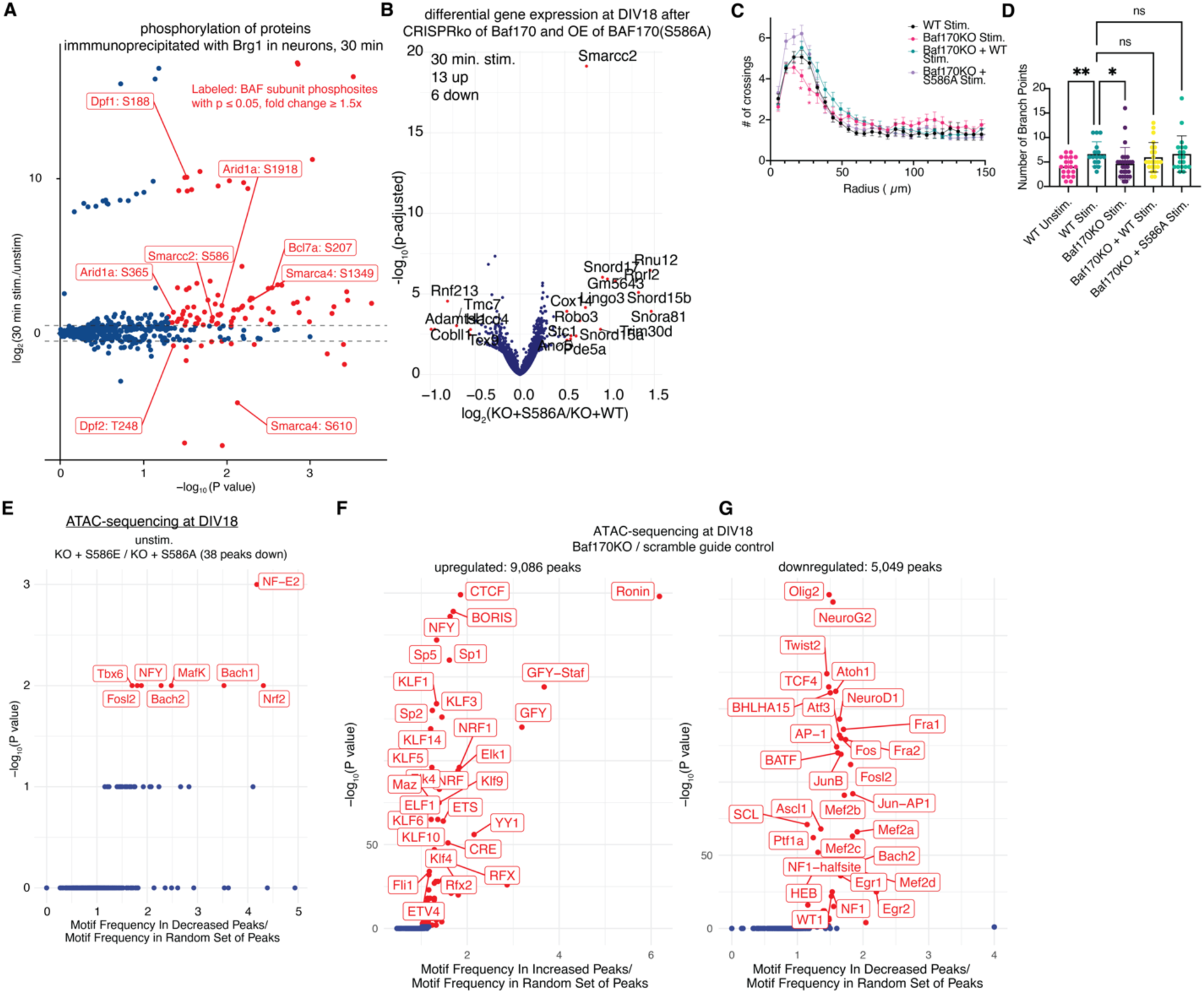
Effects of activity-dependent phosphorylation of BAF on chromatin accessibility. **A.** Differential phosphosites (norm. respective protein levels) detected after 30 min. stimulation of cortical neurons; P-values computed by two-sided Wald test, differential sites were classified as p_adj_≤0.05 and |log_2_(change)| ≥ 1.5. **B.** Volcano plot of changes after 30 min. stimulation of DIV18 cortical neurons between those that had Baf170KO+Baf170(S586A) and those with Baf170KO+ Baf170(wild-type, WT) added back. n=2- 3 biological replicates. P-values computed by two-sided Wald test, adjusted for multiple hypotheses by Benjamini-Hochberg; differential (labeled in red) peaks were classified as p_adj_≤0.05 and |log_2_(change)|≥ 0.5. **C.** Sholl analysis of activity-dependent dendritic arborization in cortical neurons with changes in Baf170 status; *: p ≤0.05, Students’ t-test; n=20-30 neurons in each condition. **D.** Number of branch points detected in Sholl analysis from **C. E.** Enrichment of transcription factor motifs in decreased (chromatin accessibility peaks in DIV18 neurons that change between Baf170 KO + S586 status; P-values computed by two-sided Fishers’ exact test and q-value adjusted by Benjamini-Hochberg, labeled in red are q < 0.05; n=2 biological replicates. **F., G.** Enrichment of transcription factor motifs in differential chromatin accessibility peaks in DIV18 neurons that are increased (**F**) or decreased (**G**) between Baf170 KO or scramble guide control (wild-type) status; P-values computed by two-sided Fishers’ exact test and q-value adjusted by Benjamini-Hochberg, labeled in red are q < 0.05; n=2 biological replicates.

**Supplemental Figure 5.**
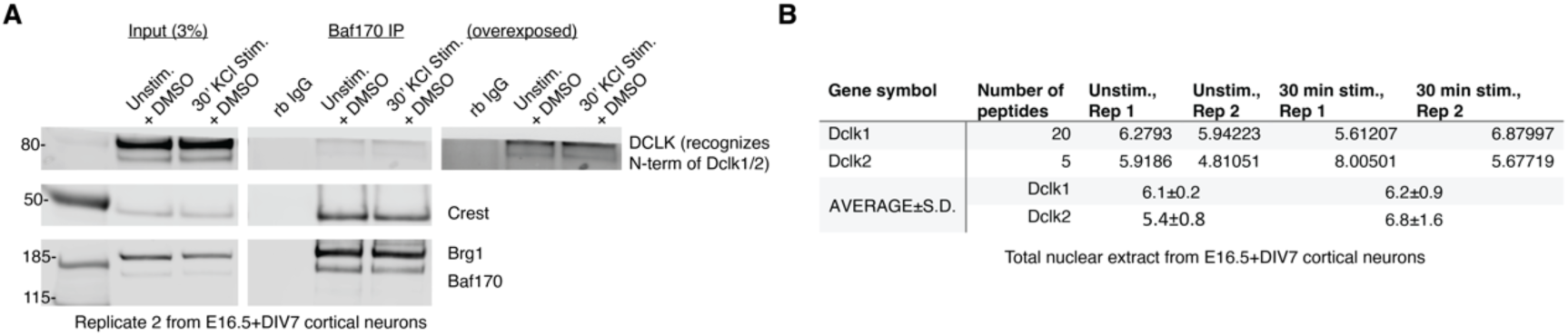
Dclk and BAF. **A.** Second replicate of Baf170 co-immunoprecipitation of Dclk1/2 in cortical neurons. **B.** Total peptides and normalized relative abundance measurements of Dclk1 and Dclk2 in nuclear extract MS.

**Supplemental Figure 6.** **A.** Full scans of Western blots.

**Supplemental Table 1.** All protein and phosphopeptides and relative abundance measurements in neuronal nuclear extract MS.

**Supplemental Table 2.** All protein and phosphopeptides and relative abundance measurements in neuronal Brg1-IP-MS.

