## Supplemental Figure 6 for "Synaptic Activity Causes Minute-scale Changes in BAF Complex Composition and Function"

**A.** Full scans of Western blots.

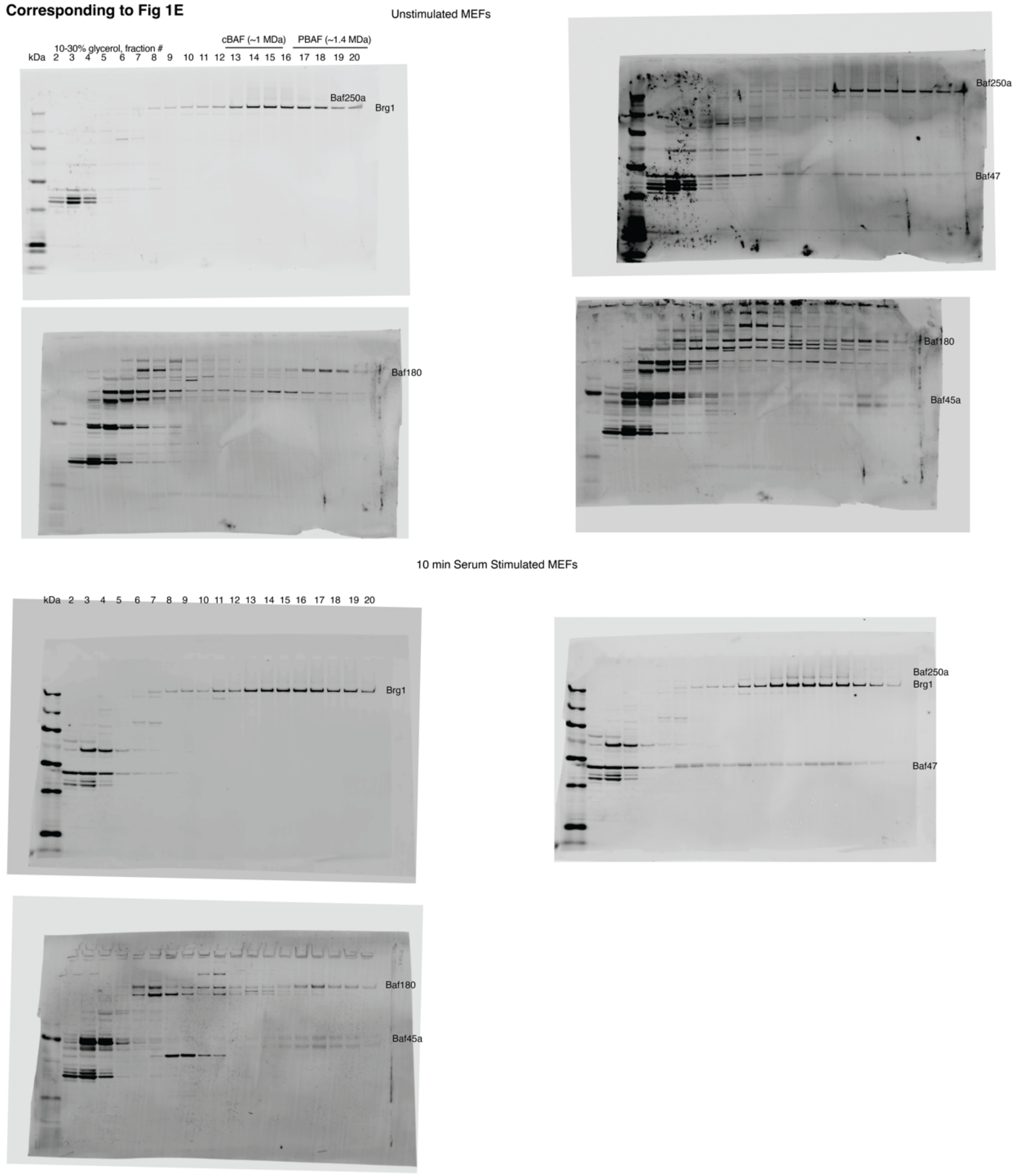

30 min Serum Stimulated MEFs

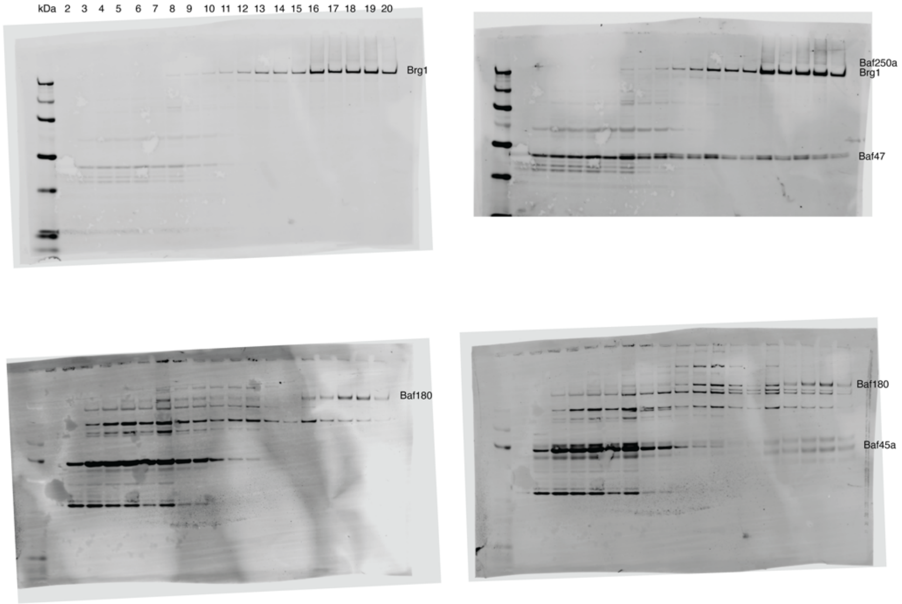


**Corresponding to Fig 1F**

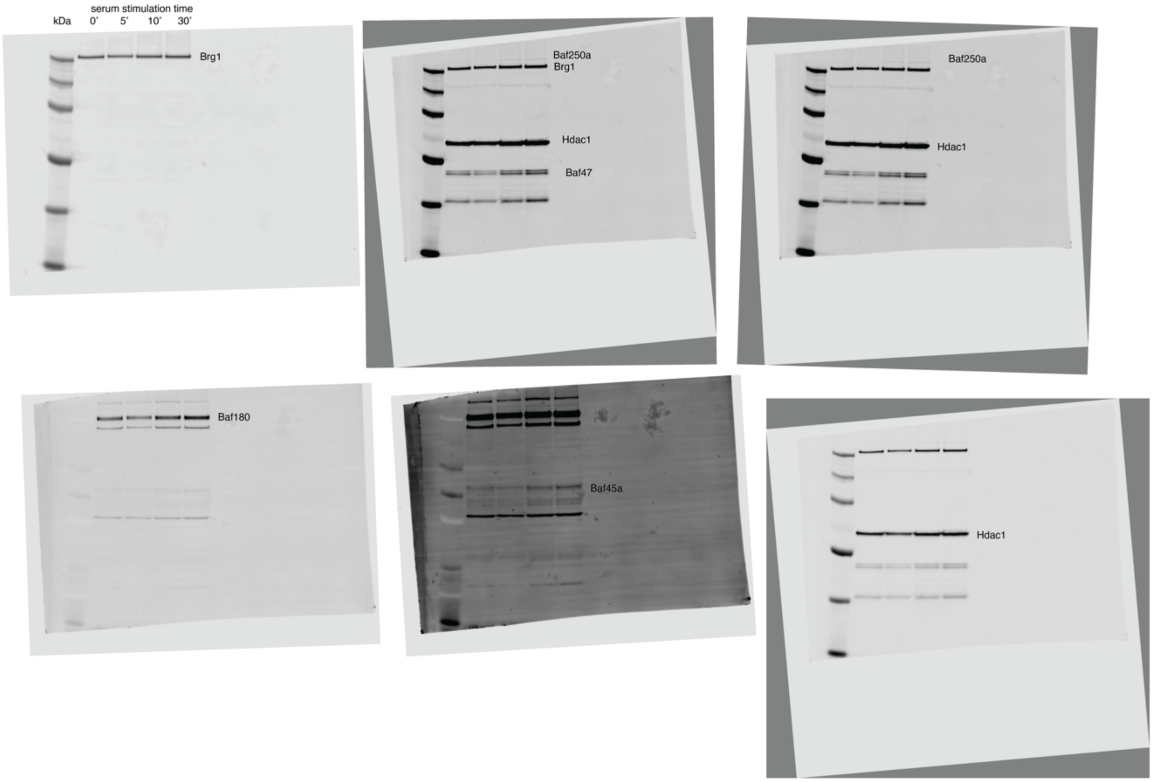

**Corresponding to Fig 3E**

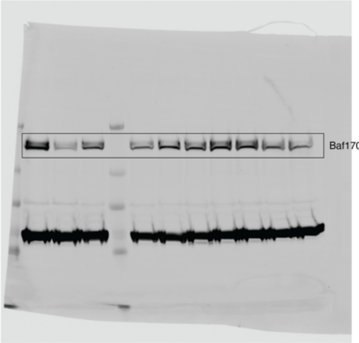


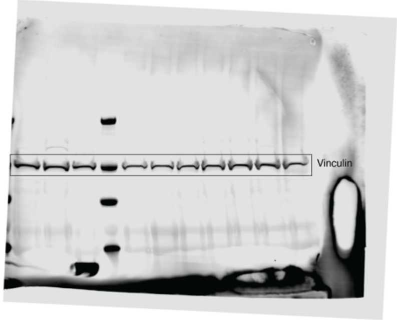


**Corresponding to Fig 4C**

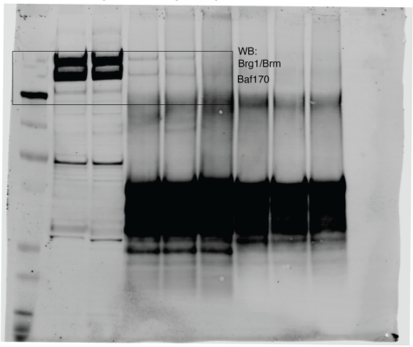


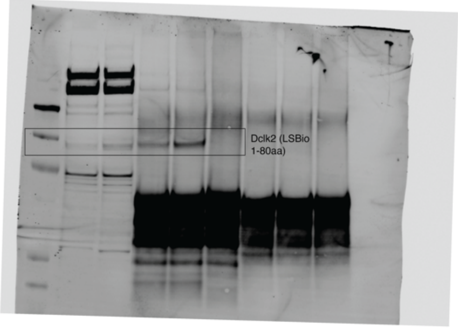


**Corresponding to Fig 4D**

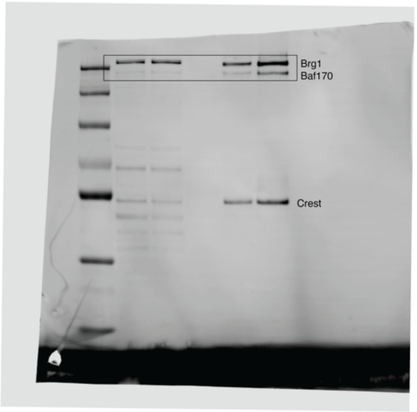


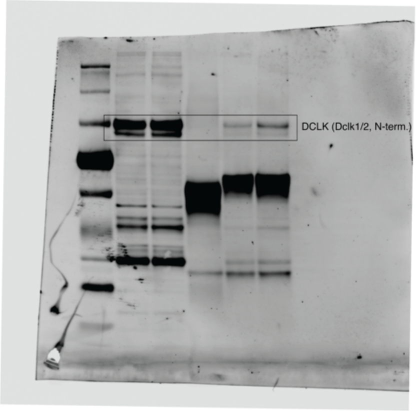

**Corresponding to Fig 4E**

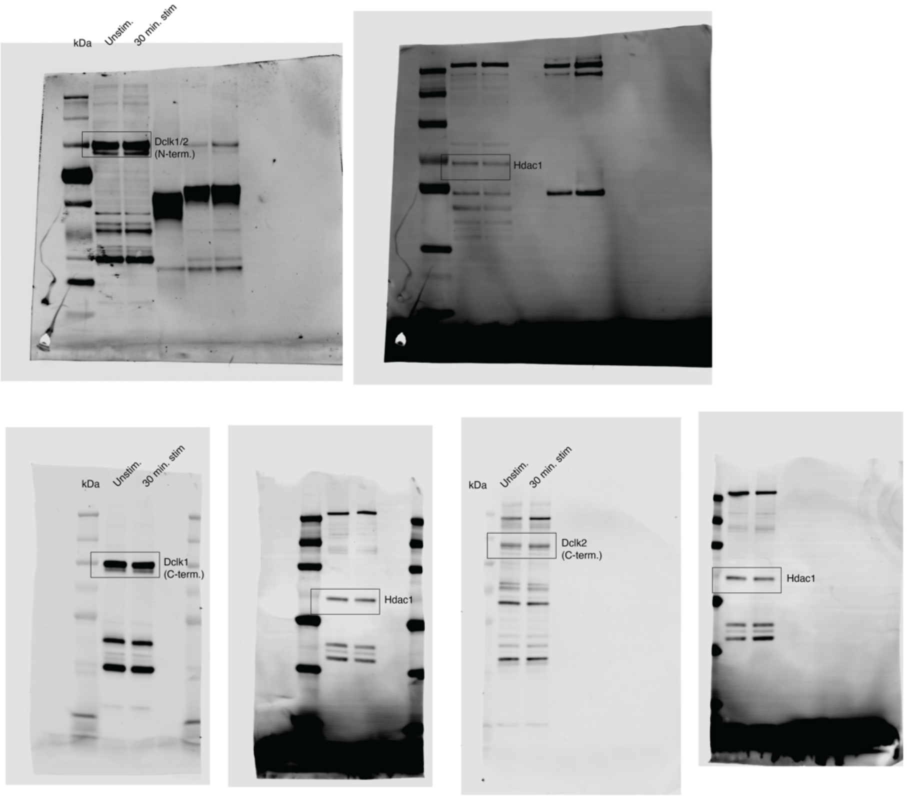


**Corresponding to Fig 4F**

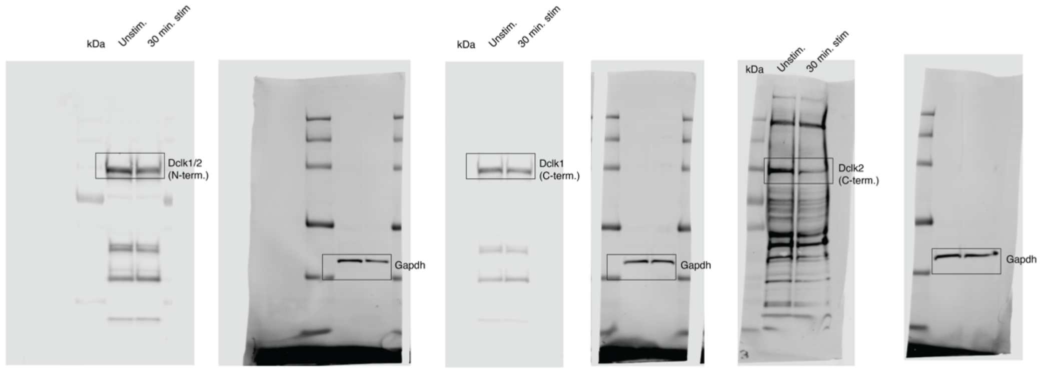

### Corresponding to Supplemental Fig 5A

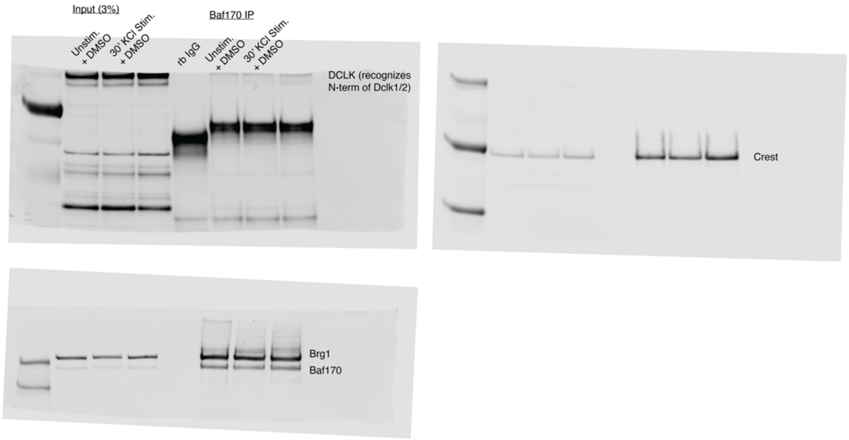
